# Interclonal cooperation and suppression shape early Ras-driven tumour growth

**DOI:** 10.64898/2026.02.19.706761

**Authors:** José Teles-Reis, Caroline Dillard, Marina Gonçalves Antunes, Ashish Jain, Paula Ruiz-Duran, Min Deng, Dan Liu, Clement Gaudin, Sigve Nakken, Amrinder Singh, Michael E Baumgartner, Anna M. Dahlström, Vilde Reinertsen, Tor Erik Rusten

## Abstract

Cancer is generally thought to be caused by expansion of a single mutant cell. However, analyses of human early lesions show that tumours can originate from several genetically distinct cell populations^1–5^. How neighbouring mutant clones interact to shape tumourigenesis, and which driver genes mediate these effects is largely unexplored. Here, we use an *in vivo* mosaic *Drosophila* epithelial model to systematically test interclonal interactions during early Ras-driven tumourigenesis. We screened 88 recurrent *RAS* co-mutated driver genes in human carcinomas for their ability to modify Ras-clone growth when disrupted in neighbouring epithelial clones. This uncovered two opposing classes of interactions: Interclonal cooperativity, where neighbouring mutant clones promote the overgrowth of Ras mutant tumours, and interclonal suppression, in which neighbours restrain Ras tumours, unexpectedly improving host survival. The strongest suppressive modifiers included canonical cell competition regulators, including *Myc*, *archipelago* (ago/*FBXW7*), and *taiman* (tai/*NCOA1-3*). In contrast, the strongest cooperative modifiers were disruptions of *XNP/ATRX and* SWI/SNF chromatin remodelling subunits (including *Osa*/*ARID1A*, *Bap170*/*ARID2*, *Polybromo*/*PBRM1*, among others), which in neighbouring cells induce a wound-like inflammatory program and drive an anabolic, pro-growth state in Ras tumours. Notably, the disruption of SWI/SNF components cell autonomously within Ras tumours confers no cooperativity. We show that interclonal cooperative Ras tumour growth requires reactive oxidative species and prostaglandin synthesis in SWI/SNF-disrupted clones. Together, this study provides a catalogue and emerging principles of cooperative and suppressive interclonal interactions among recurrent *RAS* co-mutated drivers, extending the rules of oncogenic cooperation beyond cell intrinsic co-mutation.

## Introduction

Tumour initiation is often portrayed as the clonal expansion of a single cell that acquires advantageous driver mutations over time. By contrast, early human solid tumour lesions have been shown to commonly display polyclonal architectures, in which several genetically distinct clones emerge and expand in parallel^1–5^. Studies from pre-cancerous epithelia have been especially informative. In the human colon, genome sequencing analyses estimated that 40% of premalignant polyps with benign histology and 28% with dysplasia have a polyclonal origin^5^. Remarkably, even within a single oncogene, *KRAS*, analogous but distinct activating mutations appear in separate clones^3^. Whether such neighbouring clones, with distinct oncogene (Onc) and tumour suppressor gene (TSG) mutations, merely co-occur, or instead cooperate, or repress each other to shape growth and selection during the earliest stages of tumour evolution, remains poorly defined.

Activating mutations in the *RAS* gene family are among the most common and earliest genetic alterations in human carcinomas^6,7^. Yet, seminal studies have shown that *RAS* mutations alone are insufficient to drive cancer development in most tissues^8^ and may even compromise cell viability^9^. Mechanistic insights into this paradox have revealed that oncogenic RAS can trigger anti-tumourigenic programs such as oncogene-induced senescence^9–11^, and that *RAS-*mutant cells can be actively eliminated from epithelial tissues through apical extrusion mediated by neighbouring cells^12^.

Early tumour development of RAS-driven lesions therefore appears to depend on a combination of cell-intrinsic and extrinsic cooperating events. Indeed, several cell-autonomous mutations, such as *TP53*^13,9^, *LKB1*^14^, *TGFBR2*^15^, *RASSF1A*^16^, *SCRIB*^17^, have been shown to synergize with oncogenic RAS to promote tumour progression. In addition, non-cell autonomous phenomena such as tissue inflammation, injury, and diet, have been shown to serve as key factors that can accelerate RAS-driven tumour growth^18–25^. Notably, despite RAS mutations representing an early transformation event, RAS-mutant clones have been found to coexist with other genetically distinct clones in early colorectal lesions, rather than dominating the entire tumour^3,5^, raising the possibility that Ras-driven tumour progression is shaped by interactions with neighbouring tumour clones.

Several studies have demonstrated that the behaviour of individual tumour clones can be shaped by their surrounding clonal landscape^26–30^. Notably, work in *Drosophila* models has been instrumental in identifying interclonal interaction behaviours and dissecting the underlying mechanisms. Wu *et al.* discovered that clones expressing oncogenic Ras (*Ras^V12^*) in the larval eye-antennal disc (EAD) exhibited limited growth on their own. However, when adjacent to polarity deficient *scribble* (scrib) mutant clones, *Ras^V12^* clones underwent massive overgrowth and invaded the neighbouring central nervous system^26^. This behaviour was explained by Upd/IL-6 cytokine production by *scrib* mutant cells via activation of the JNK pathway, and activation of pro-growth and invasion JAK-STAT pathway in *Ras^V12^* cells^26^. In another study, Enomoto *et al.* found that neighbouring *Ras^V12^* and *Src* overexpressing cells (*Src^OE^*) mutually stimulate each other’s invasion potential. *Ras^V12^* clones promoted *Src^OE^* invasion through Delta-Notch signalling, while *Src^OE^* induced *Ras^V12^* invasion by activating the JAK-STAT pathway through Upd/IL-6 release^27^.

In mice studies, Cleary *et al.* showed in a Wnt1-driven mouse mammary tumour model, that genetically distinct subclones arise and cooperate to sustain tumour growth. Spontaneously arising *Hras* mutant subclones, which express low levels of Wnt1, depend on neighbouring Wnt1-expressing *Hras* wildtype tumour cells. Neither subclone alone is sufficient for tumour generation *in vivo*^28^. Furthermore, Marusyk *et al.* showed in a xenograft breast cancer model that an IL-11 overexpressing subclone could promote tumour growth through non-autonomously stimulating expansion of neighbouring subclones^29^. In another study, Amara *et al.* showed that only a multiclonal mixture of subclones isolated from a patient-derived ovarian carcinoma line could form solid peritoneal metastases when injected in mice. These metastases were dominated by an ERBB2-amplified subclone, which depended on the ERBB2 amphiregulin ligand produced by other subclones, revealing an interclonal cooperativity behaviour^30^.

Whether interclonal interactions can suppress a neighbouring tumour clone is much less well characterised. However, a recent study suggests that *Notch1* deficient clones, which are highly pervasive in the epithelia of healthy individuals^31,32^, may exert an interclonal suppressive effect. Colom *et al.* showed that in a mouse model where oesophageal tumours are induced by carcinogen treatment in drinking water, the generation of Notch signalling deficient clones significantly suppressed the growth of neighbouring nascent tumours^33^.

Despite growing evidence of interclonal interactions in cancer, no systematic effort has been made to identify the role of frequently mutated oncogenes and tumour suppressors in driving these interactions. In this study, using EyaHOST, a recently developed *Drosophila* tool that enables the individual genetic manipulation of two distinct epithelial cell populations *in vivo*^34^, we have performed a genetic screen for clinically relevant Onc and TSGs that promote interclonal interaction behaviours with *Ras^V12^*clones. We have identified five different TSGs that when disrupted in neighbouring cells can non-autonomously promote the overgrowth of *Ras^V12^* tumours. Conversely, we have found that the misexpression of two TSGs and three Onc can interclonally suppress the growth of *Ras^V12^,* resulting in increased survival of these animals. Several genes identified in our screen that interclonally promote *Ras^V12^* tumour growth belong to the SWItch/Sucrose Non-Fermentable (SWI/SNF) complex, a chromatin-remodelling complex that is recurrently mutated across human cancers^35^. Disruption of SWI/SNF induces a wound-like inflammatory state characterised by elevated caspase activity, JAK-STAT and JNK signalling, reactive oxygen species (ROS), and DNA damage. In response, *Ras^V12^* clones undergo an anabolic shift marked by increased mechanistic Target of rapamycin (mTOR) activity and protein translation. Neighbouring SWI/SNF-disrupted clones robustly promote *Ras^V12^* tumour overgrowth, a phenotype that was found to depend on ROS and prostaglandin synthesis within SWI/SNF-deficient cells.

This work expands the repertoire of the mutational drivers capable of inducing interclonal interactions and reveals new mechanisms through which interclonal crosstalk contributes to tumour progression.

## Results

### A reverse genetic screen identifies clinically relevant oncogenes and tumour suppressors that promote interclonal interactions

To identify oncogenes and tumour suppressor genes that can non-autonomously alter the behaviour of *Ras^V12^* expressing cells, we established an assay to study interclonal interactions using EyaHOST^34^ (Fig. 1A). This recently developed genetic tool enables the manipulation of host tissues independently of tumour initiation and was designed to study tumour-host interactions. We repurposed this tool to investigate interclonal interactions by using a version of EyaHOST that allows genetic manipulation of epithelial cells adjacent to *Ras^V12^* clones^34^. In these neighbouring cells, we aimed to introduce additional tumourigenic alterations by knocking down TSGs or overexpressing Onc of interest, and assess how *Ras^V12^* cells respond when confronted with these genetically altered clones.

**Fig. 1.**
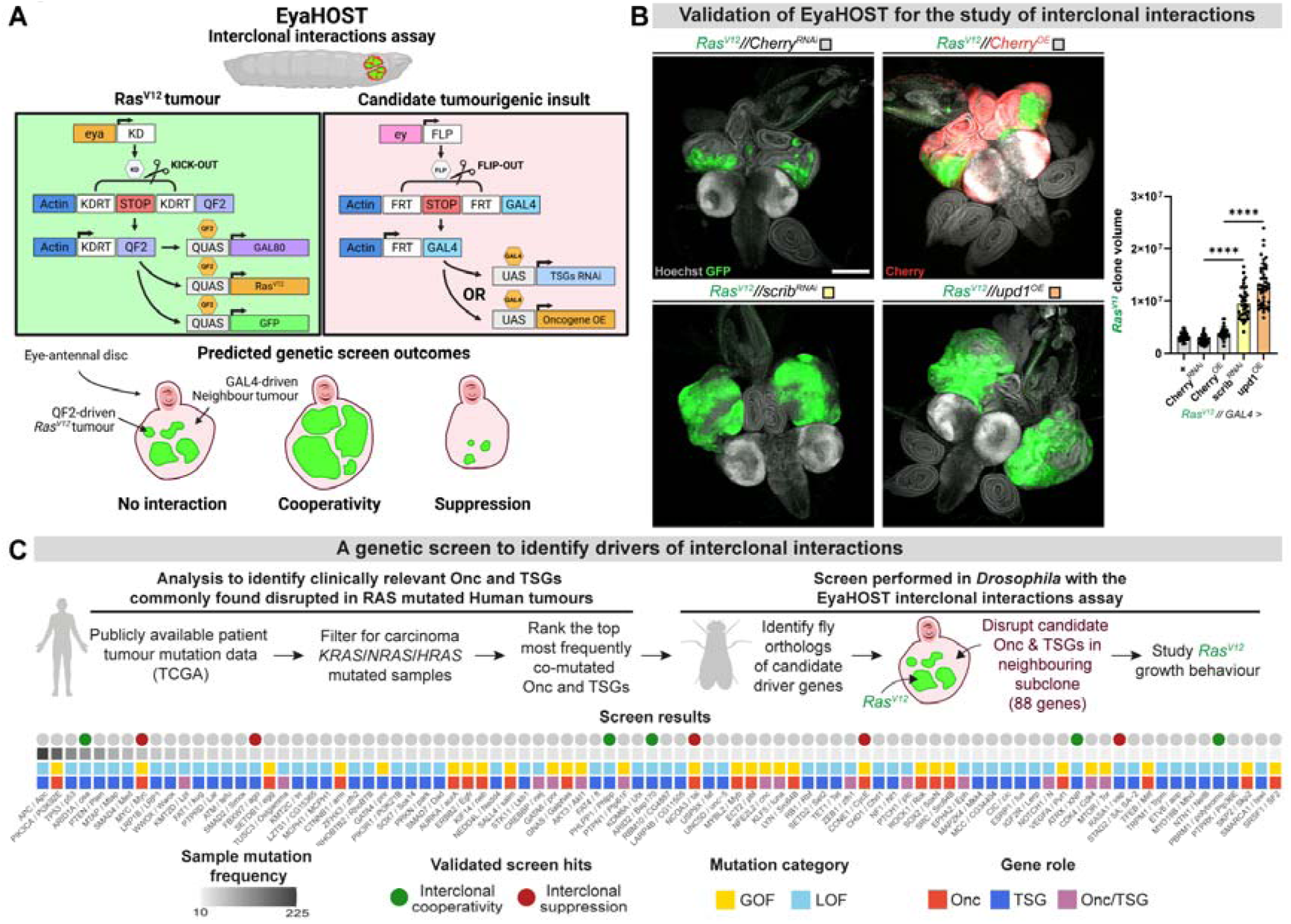
A genetic screen for the identification of clinically relevant Onc/TSGs promoting interclonal interactions. (A) Schematic illustrating the genetic strategy used in the EyaHOST interclonal interactions assay. *Ras^V12^*tumour clones are induced in the EAD using the eya-driven KD/KDRT recombination system. This “kick-out” approach removes a STOP cassette, triggering constitutive activation of the QF2-QUAS system in only a few cells of the EAD, which drives expression of Ras^V12^, GFP, and GAL80. In parallel, *ey*-driven expression of FLP recombinase mediates “flip-out” of an FRT-flanked STOP cassette, leading to GAL4 expression in the whole EAD. Because QF2-QUAS and GAL4-UAS are orthogonal systems, and because GAL80 inhibits GAL4 activity within QF2^+^ clones, the two populations can be manipulated independently. In the GAL4^+^ neighbouring cells we introduce a second tumour clone by RNAi or overexpression of candidate TSGs or Onc, respectively. The assay then aims to study the growth behaviour of *Ras^V12^*clones when challenged by a neighbouring tumour clone. We predicted three screen outcomes, which were of no interaction (no changes to *Ras^V12^* growth), cooperativity (increased *Ras^V12^* growth), or suppression (reduced *Ras^V12^* growth). (B) Validation of the EyaHOST system for the study of interclonal interactions. Previous study has shown that *scrib* loss and *upd* upregulation could non-autonomously promote *Ras^V12^*overgrowth^26^. To validate EyaHOST, we compared *Ras^V12^* clone behaviour in cephalic complexes of controls *Ras^V12^//+* (n=33), *Ras^V12^//Cherry^RNAi^* (n=46) and *Ras^V12^//Cherry^OE^* (n=29) to *Ras^V12^//scrib^RNAi^* (n=40) and *Ras^V12^//upd^OE^* (n=43). Representative confocal images show Hoechst-stained nuclei (grey) and *Ras^V12^* clones marked with GFP (green). The Cherry pattern (red) in *Ras^V12^//Cherry^OE^* confirms the system’s capacity for non-autonomous manipulation. *Ras^V12^*clone volume (µm^3^) of distinct cephalic complexes was quantified, and statistical significance was determined with one-way ANOVA and Tukey’s multiple comparison test. Scale bar = 400 µm. (C) Workflow of the interclonal interactions genetic screen. To identify clinically relevant drivers of interclonal interactions, we analyzed publicly available TCGA datasets to find Onc and TSGs frequently co-mutated in KRAS/NRAS/HRAS-mutant human carcinomas. We then identified *Drosophila melanogaster* orthologs and selected transgenic lines to disrupt 88 candidate genes in the *Ras^V12^*neighbouring cells with EyaHOST. Candidate genes were prioritized based on co-mutation frequency and availability of fly lines. The bottom panel shows genes ranked by co-mutation frequency with *Ras^V12^*, annotated with both human and fly gene symbols. Validated hits are color-coded by screen outcome: green for interclonal cooperativity, red for interclonal suppression. Each gene is also labelled by mutation type (GOF or LOF) and functional category (Onc, TSG, or both).

EyaHOST allows the independent genetic manipulation of two epithelial cell populations in the *Drosophila* larval EAD through a dual-recombinase and dual-binary expression system approach^34^. Briefly, for *Ras^V12^* tumour generation, the *eyes absent* (*eya*) enhancer-driven KD recombinase “kicks-out” a *KDRT-STOP-KDRT* cassette in only a few random cells of the EAD, generating clones which have constitutive expression of the QF2 transcription factor. This then activates *QUAS-Ras^V12^* and *QUAS-GAL80* transcription, together with GFP labelling. In parallel, an alternative recombinase system FLP/FRT is induced in the whole EAD through the *eyeless* (*ey*) enhancer, which “flips-out” an *FRT-STOP-FRT* cassette and allows constitutive activity of the GAL4-UAS system in the whole tissue. The GAL4 inhibitor, GAL80, expression inside the QF2^+^ clones represses any GAL4 activity, resulting in two mutually exclusive epithelial cell populations where GAL4 can be used to manipulate any gene of choice (Fig. 1A).

Seminal work by Wu *et al.* revealed that *scrib^-^* clones can promote the overgrowth of neighbouring *Ras^V12^* tumours through the secretion and paracrine signalling by Upd/IL6^26^. To validate the EyaHOST interclonal interactions assay (Fig. 1A), we sought to recapitulate these previous findings^26^. As expected, we observed no difference between the different controls tested *Ras^V12^//+* (the // symbol denotes manipulation in two neighbouring epithelial compartments in the EAD), *Ras^V12^//Cherry^RNAi^*, and *Ras^V12^//Cherry^OE^* (Fig. 1B). However, upon either downregulation of *scrib* or overexpression of *upd1* in neighbouring cells, we observed a strong increase in *Ras^V12^* growth (Fig. 1B), which validated our assay approach.

To enhance the clinical relevance of our screen, we focused on genes frequently altered in human tumours. We hypothesized that genes capable of promoting interclonal interactions with *Ras^V12^* would appear frequently co-mutated with *RAS* in patient tumour samples. Moreover, because we are modelling epithelial tumours using the *Drosophila* larva EAD, we restricted our analysis to genes altered in human carcinomas. Based on these criteria, we ranked the most frequently RAS co-mutated Onc and TSGs present in publicly available patient tumour datasets (TCGA) and determined their corresponding *Drosophila* orthologues (Fig. 1C). Transgenic lines to perform either RNA interference (RNAi) for TSGs and UAS-mediated overexpression lines for Onc were obtained, resulting in a total of 88 candidate genes. We anticipated there would be three screen outcomes: the second neighbouring tumour population would not affect *Ras^V12^* growth (no interaction), it could induce the growth of *Ras^V12^* (interclonal cooperativity) or suppress the expansion of *Ras^V12^* (interclonal suppression) (Fig. 1A). The screen was carried out by two independent scorers in a blinded manner, by visual qualitative assessment of GFP-labelled *Ras^V12^* tumour size under a fluorescent stereoscope.

Strikingly, knockdown of five distinct TSGs (*osa*, *Bap170*, *polybromo*, *XNP*, and *Phlpp*) in neighbouring cells led to an apparent non-cell autonomous enhancement of *Ras^V12^* tumour overgrowth (Fig. 1C). In contrast, misexpression of two TSGs (*vap* and *ago*) and three Onc (*tai*, *myc*, and *CycE*) in neighbouring cells seemed to result in diminished *Ras^V12^* tumour growth (Fig. 1C). Thus, the phenotypic categories observed fit the initial predicted outcomes of our screen. Notably, most Onc or TSGs had no observable effect on neighbouring *Ras^V12^* growth, whereas interclonal cooperativity and suppression were observed only with a few genes, indicating specificity of interactions.

### Non-autonomous *Ras^V12^*overgrowth is triggered by the loss of five distinct TSG in neighbouring epithelium

To validate our screen results, we dissected and compared EADs from control (*Ras^V12^//Cherry^RNAi^*) to EADs with knockdown in *Ras^V12^* neighbouring cells for the five different TSG hits (*Ras^V12^//osa^RNAi^, Ras^V12^//Bap170^RNAi^, Ras^V12^//polybromo^RNAi^, Ras^V12^//XNP^RNAi^, and Ras^V12^//Phlpp^RNAi^*) (Fig. 2A).

**Fig. 2.**
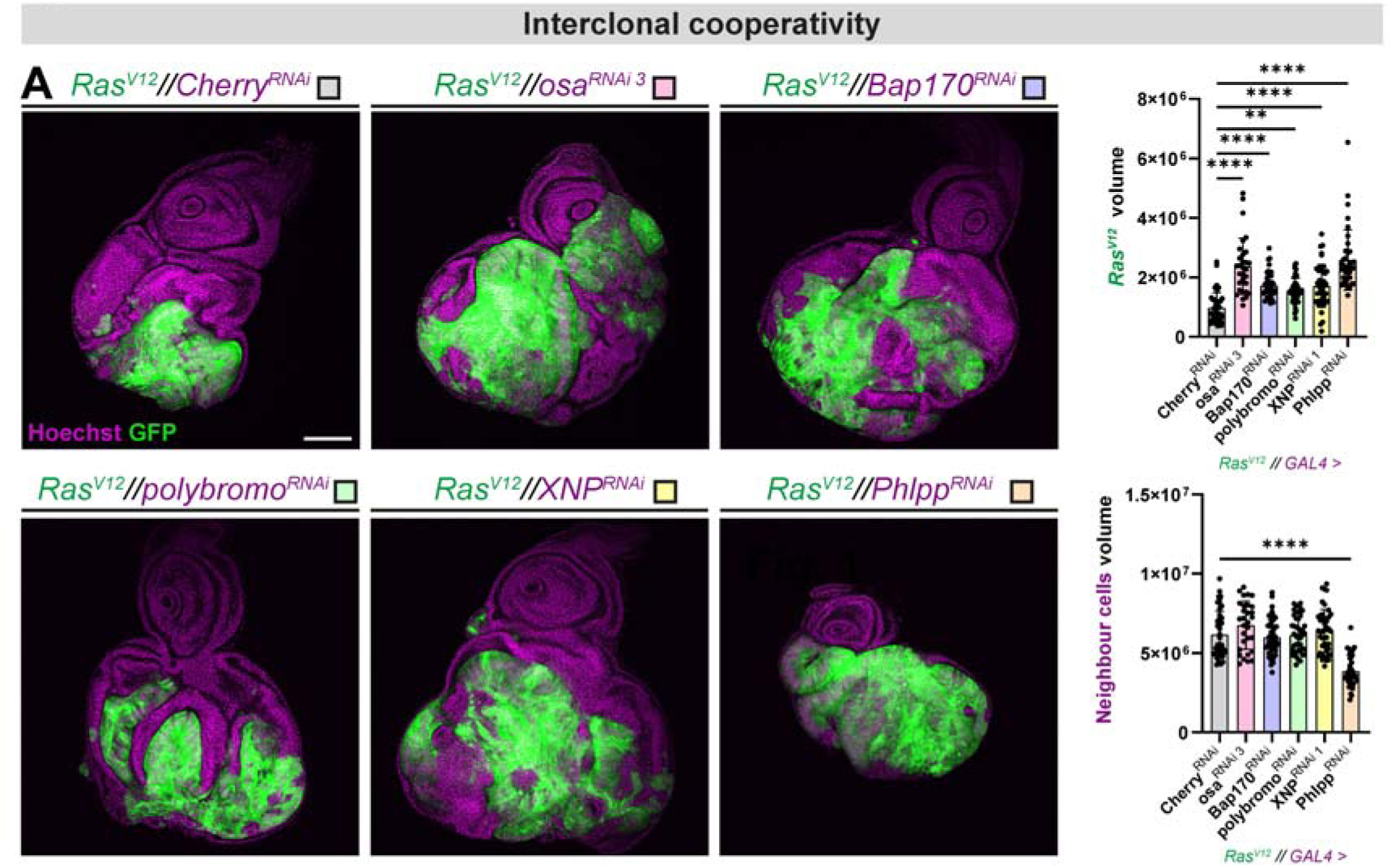
Disruption of 5 different TSGs in neighbouring cells promotes *Ras^V12^* overgrowth. (A) Representative confocal images of EADs showing control *Ras^V12^//Cherry^RNAi^* (n=42) and the effect of knocking down in *Ras^V12^* neighbouring cells five different TSGs *osa* (n=30), *Bap170* (n=45), *polybromo* (n=42), *XNP* (n=44), *and Phlpp* (n=37) which were identified in the screen as promoters of interclonal cooperativity. In each condition, *Ras^V12^* cells are labelled with GFP (green), and nuclei are stained with Hoechst (magenta). Quantification of *Ras^V12^* clone volume (top right) and neighbouring tissue volume (bottom right) of distinct EADs is shown (µm^3^). Statistical significance was assessed with Kruskal-Wallis and Dunn’s multiple comparison test. Scale bar = 100 µm.

Among the initial screen hits, *osa*, *Brahma-associated protein 170kD* (Bap170), and *polybromo*, all encode components of SWI/SNF, an evolutionarily conserved ATP-dependent chromatin remodelling multi-protein complex involved in nucleosome mobilization^35^. SWI/SNF regulates transcription by remodelling chromatin and modulating the recruitment of transcriptional regulators at regulatory elements. In addition, it plays a role in the DNA damage response by facilitating the access of repair factors to damaged sites^35^. Non-autonomous disruption of either of these genes individually resulted in increased *Ras^V12^* overgrowth, while not affecting the volume of the neighbouring subclone with SWI/SNF component knockdown (Fig. 2A). We verified that we can specifically promote downregulation of SWI/SNF subunits in *Ras^V12^*surrounding cells (Suppl. fig. 2A). The independent identification of three genes from the same complex in our genetic screen (Fig. 1C, 2A) strongly suggested that disrupted SWI/SNF activity can promote the non-autonomous induction of *Ras^V12^* overgrowth.

Furthermore, we identified *XNP*, a heterochromatin regulator orthologous to human *ATRX*, which, similar to SWI/SNF subunit Brahma, contains a SNF2 ATPase domain essential for nucleosome remodelling. As a chromatin remodeller, ATRX has been shown to modulate transcription and DNA damage repair^36,37^. Indeed, upon downregulation of *XNP* in neighbouring cells by two independent RNAi lines, we observed an increase in the growth of *Ras^V12^* (Fig. 2A, Suppl. fig 1A). *Drosophila* XNP contains high homology to the human ortholog ATRX SNF2 domain. However, XNP is missing the ADD domain from ATRX, which is important for heterochromatin binding. In flies, this domain is instead encoded by another gene, *ADD1*, which has been shown to interact with *XNP*^37^. *ATRX* is believed to have undergone fission in insects during evolution to generate *XNP* and *ADD1*^38^. To test whether the non-autonomous growth effect of *XNP* loss was also phenocopied by *ADD1*, we similarly downregulated *ADD1* in *Ras^V12^* surrounding cells with two independent RNAis, for which we observed no effect (Suppl. fig 1A). Thus, the non-autonomous effect upon *XNP* downregulation possibly depends on other binding partners. Indeed, in polytene chromosomes, XNP has been shown to overlap or to bind different regions than ADD1^37^. Furthermore, it has been described that a short XNP isoform does not contain an ADD1 binding site and instead preferentially localizes to euchromatin regions^37^.

We additionally identified *Phlpp*, a conserved serine/threonine phosphatase, shown to control cell polarity^39^ and to negatively regulate AKT activity by dephosphorylation, thereby promoting apoptosis^40^. Upon knockdown of *Phlpp* in surrounding cells we observed an increase in *Ras^V12^* clone volume (Fig. 2A). However, we were surprised to observe a strong reduction in tissue size in the neighbouring compartment (Fig. 2A). Given that Phlpp is known to negatively regulate AKT signalling, its knockdown was expected to enhance AKT activity and promote cell survival. In contrast, we observed increased cell death, as indicated by an increase in pyknotic nuclei (data not shown). It is possible that tumour suppressive mechanisms are activated in response to *Phlpp* knockdown, such as enhanced JNK signalling, which may account for the increased cell death and tissue reduction, which potentially can explain the non-autonomous *Ras^V12^* overgrowth.

Together, these results validate our genetic screening approach and reveal multiple chromatin remodelling factors that, when disrupted in the surrounding tissue, promote non-autonomous *Ras^V12^* growth, uncovering novel mutational drivers of interclonal interactions.

### Perturbation of specific Onc and TSG in surrounding cells suppresses *Ras^V12^*-driven growth and promotes animal survival

Evidence that neighbouring clones can outcompete other surrounding tumour cells remains limited. However, we hypothesized that disruption of certain Onc and TSGs could render neighbouring cells more fit, thereby leading to the suppression of nascent *Ras^V12^* tumours.

Work in *Drosophila* has been instrumental in establishing the concept of cell competition, a process in which insulted unfit cells, rendered so by gene mutations or cellular stress, can be actively eliminated by fitter neighbouring cells^41^. This phenomenon is critical for tissue homeostasis, but can be hijacked by cancer cells, whereby certain oncogenic alterations can induce the opposite behaviour. This process is termed supercompetition and is observed when the insulted cell acquires the capacity to induce the elimination of surrounding cells^41^. Supercompetitive behaviour was first described in cells expressing relatively higher levels of *myc* to its neighbours^42^.

Strikingly, we observed that *myc* upregulation in cells surrounding *Ras^V12^*clones led to a marked reduction in *Ras^V12^* clone volume (Fig. 3A). This phenomenon is reminiscent of supercompetition, in which *myc^OE^* cells eliminate neighbouring wildtype cells. In our experiments, we find that this competitive behaviour extends to the non-autonomous elimination of nascent *Ras^V12^* tumours (Fig. 3A).

**Fig. 3.**
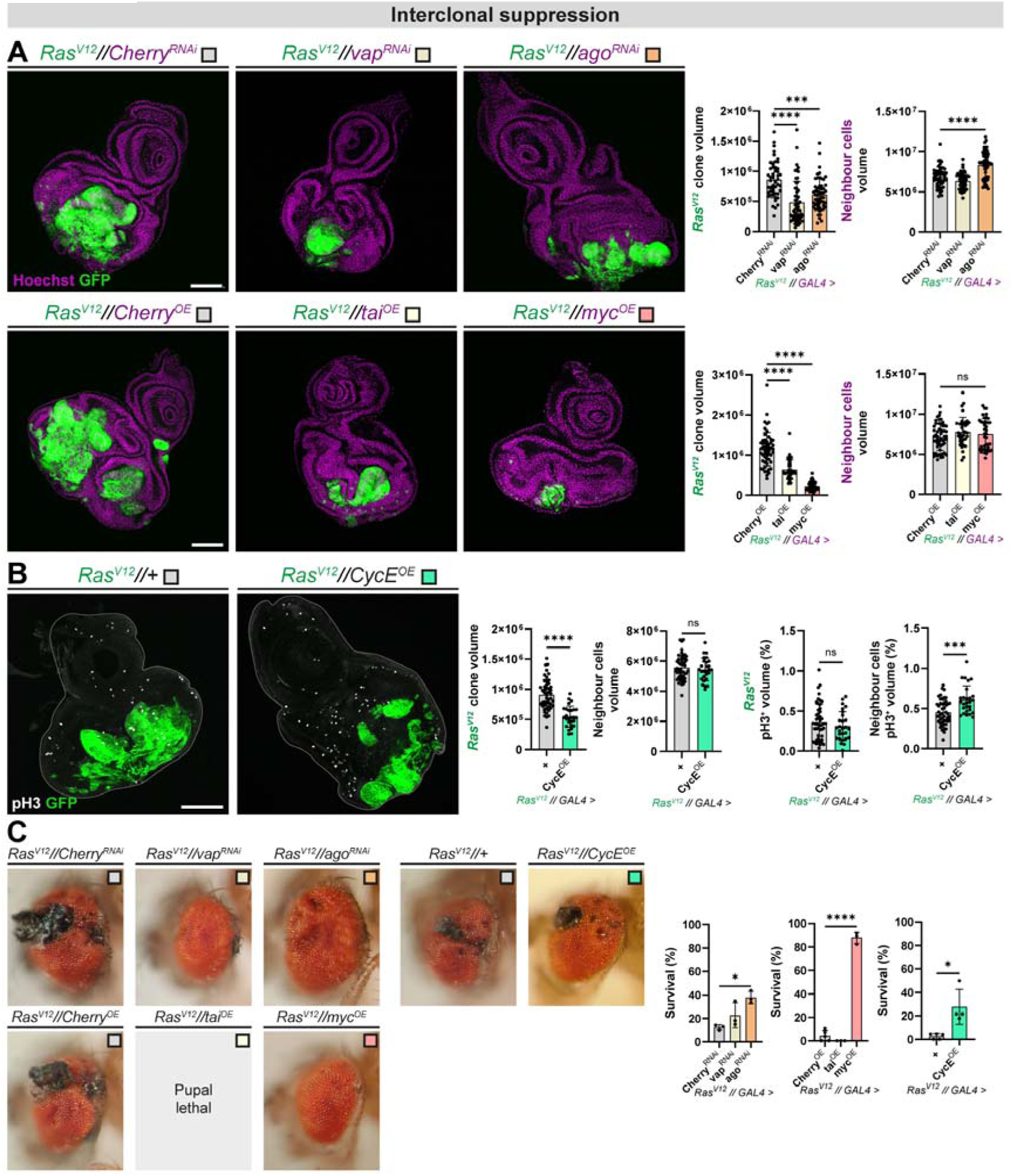
Misexpression of 3 distinct Onc and 2 TSGs in neighbouring cells suppresses *Ras^V12^* tumour growth and rescues animal lethality. (A,B) Representative confocal images of EADs showing control (*Ras^V12^//Cherry^RNAi^*, n=48; *Ras^V12^//Cherry^OE^*, n=55; and *Ras^V12^//*+, n=48) and the result of manipulating in neighbouring cells the gene hits found in the screen to promote interclonal suppression. Two tumour suppressor genes (*Ras^V12^//vap^RNAi^*, n=55; *Ras^V12^//ago^RNAi^*, n=54) were knocked down, and three oncogenes (*Ras^V12^//tai^OE^*, n=37; *Ras^V12^//myc^OE^*, n=40; *Ras^V12^//CycE^OE^*, n=27) were overexpressed non-autonomously. In each condition, *Ras^V12^* cells are labelled with GFP (green), and nuclei are stained with Hoechst (magenta). Quantification of *Ras^V12^* clone volume and neighbouring tissue volume of distinct EADs are shown (µm^3^). Additionally, pH3-positive volume fraction (%) was quantified inside *Ras^V12^* clones and in the neighbouring population in control and upon *CycE^OE^*. Statistical analyses were performed with Kruskal–Wallis and Dunn’s test (for *Ras^V12^* clone volume in RNAi and overexpression conditions, and in neighbour volumes of overexpression condition), one-way ANOVA and Tukey’s test (for RNAi neighbour volume condition), Welch’s t-test (for *Ras^V12^* clone and neighbour volumes in *CycE^OE^* comparisons), and Mann–Whitney U test (for *Ras^V12^* clone and neighbour pH3+ volume analyses). Scale bar = 100 µm. (C) Adult *Ras^V12^* survivors and eye phenotypes corresponding to the genotypes above (*Ras^V12^//Cherry^RNAi^*, n=3; *Ras^V12^//Cherry^OE^*, n=5; *Ras^V12^//*+, n=5; *Ras^V12^//vap^RNAi^*, n=3; *Ras^V12^//ago^RNAi^*, n=3; *Ras^V12^//tai^OE^*, n=3; *Ras^V12^//myc^OE^*, n=3; *Ras^V12^//CycE^OE^*, n=4). Quantifications of adult survival (Surviving adults/(Surviving adults + dead individuals) %) are shown on the right. Statistical significance was determined with one-way ANOVA and Tukey’s test (for vap, ago, tai and myc testing conditions) and with Mann–Whitney U test (for CycE overexpression experiment).

Another gene identified in our screen is *archipelago* (ago), the *Drosophila* ortholog of human *FBXW7*, an F-box protein that acts as the substrate recognition subunit of the SCF ubiquitin ligase complex. Ago has been shown to negatively regulate Myc protein levels, and its loss results in both clonal overgrowth and a reduction in neighbouring healthy tissue^43^. In line with *myc*-dependent interclonal suppression we observed, non-autonomous knockdown of *ago* also led to reduced *Ras^V12^* clone volume (Fig. 3A).

Beyond its regulation of Myc, Ago also exerts tumour suppressive activity through an independent mechanism by negatively regulating CycE levels^44^. Surprisingly, we found that upregulation of CycE in cells surrounding *Ras^V12^* tumours also induced interclonal suppression (Fig. 3B). One initial hypothesis to explain this would be that since CycE promotes G1-S phase transition, faster-cycling neighbouring cells could physically crowd and eliminate *Ras^V12^*clones. However, previous studies have shown that *CycE^OE^* cells maintain similar cell doubling time to controls due to a compensatory extension of the S phase^45^. Consistent with previous work, we did not observe a change in *CycE^OE^* compartment size (Fig. 3B). Nonetheless, we did observe an increase in cells labelled with phospho-histone H3 (pH3) in *CycE^OE^* population (Fig. 3B), which is likely explained by a prolonged mitotic phase, as previously described in cells with elevated CycE^46^. Additionally, *CycE^OE^* has been described to not change Myc levels^43^, suggesting that the observed competitive behaviour of *CycE^OE^* cells occurs via a yet unidentified mechanism.

Furthermore, the screen identified the gene *taiman* (tai), which is the *Drosophila* ortholog of the Human nuclear receptor coactivators *NCOA1-3*. Previous work has shown that cells that overexpress *tai* can also behave as supercompetitors^47^, while cells with low levels of Tai are unfit and eliminated by neighbouring healthy cells^48^. Similarly, the overexpression of this oncogene in neighbouring cells induced the suppression of *Ras^V12^*clone growth (Fig. 3A).

We additionally found in this screen the gene *vacuolar peduncle* (vap) (Fig. 3A), which is the ortholog of human *RASA1*, which encodes for the p120 Ras GTPase-activating protein (GAP) that negatively regulates the levels of Ras activation.

Furthermore, when generating *Ras^V12^* tumours with EyaHOST in the EAD we observed few larvae survive metamorphosis and hatch into adults (3-12% of individuals in control groups), while the great majority died during the pupal stage (Fig. 3C). Strikingly, we observed that several of the drivers found to promote interclonal suppression could induce an increase in adult survival. This was the case for *Ras^V12^//myc^OE^, Ras^V12^*//*ago^RNAi^*, and *Ras^V12^*//*CycE^OE^* flies (Fig. 3C). Non-autonomous *myc^OE^* induced a striking 22-fold increase in survival compared to control. Additionally, we saw an amelioration of the adult eye phenotype upon interclonal suppression (Fig. 3C), consistent with a reduction in *Ras^V12^* tumour volume (Fig. 3A, B).

Overall, these findings suggest that tuning the competitive interactions between distinct tumour clones can redirect the outcome of the clones present within the tumour tissue, potentially revealing an unexpected vulnerability of *Ras^V12^*cells to fitter neighbours.

### Disruption of core or alternative subunits of the SWI/SNF chromatin remodelling complex interclonally promotes the growth of *Ras^V12^* cells

From the identification in our screen of 3 different genes from the SWI/SNF complex, *osa*, *Bap170*, and *polybromo* (Fig. 2A), we hypothesized that disruption of this protein complex could promote the non-autonomous growth of *Ras^V12^* cells. Indeed, subunits from SWI/SNF complex have been found to act as TSGs in *Drosophila* imaginal discs^49,50^. The SWI/SNF complex exists in three forms in humans and two in *Drosophila*, distinguished by the specific variant subunits they incorporate. In *Drosophila*, these forms are known as Brahma-associated proteins (BAP) and Polybromo-associated BAP (PBAP) (Fig. 4A). The variant defining subunit identified in BAP is Osa, while the variant subunits found in PBAP are Polybromo, Bap170, and Sayp. The shared and core subunits of BAP and PBAP are Brahma (Brm), Moira (Mor), Snf5-related 1 (Snr1), Brahma-associated protein 111kD (Bap111), Brahma-associated protein 60kD (Bap60), and Brahma-associated protein 55kD (Bap55) (Fig. 4A).

**Fig. 4.**
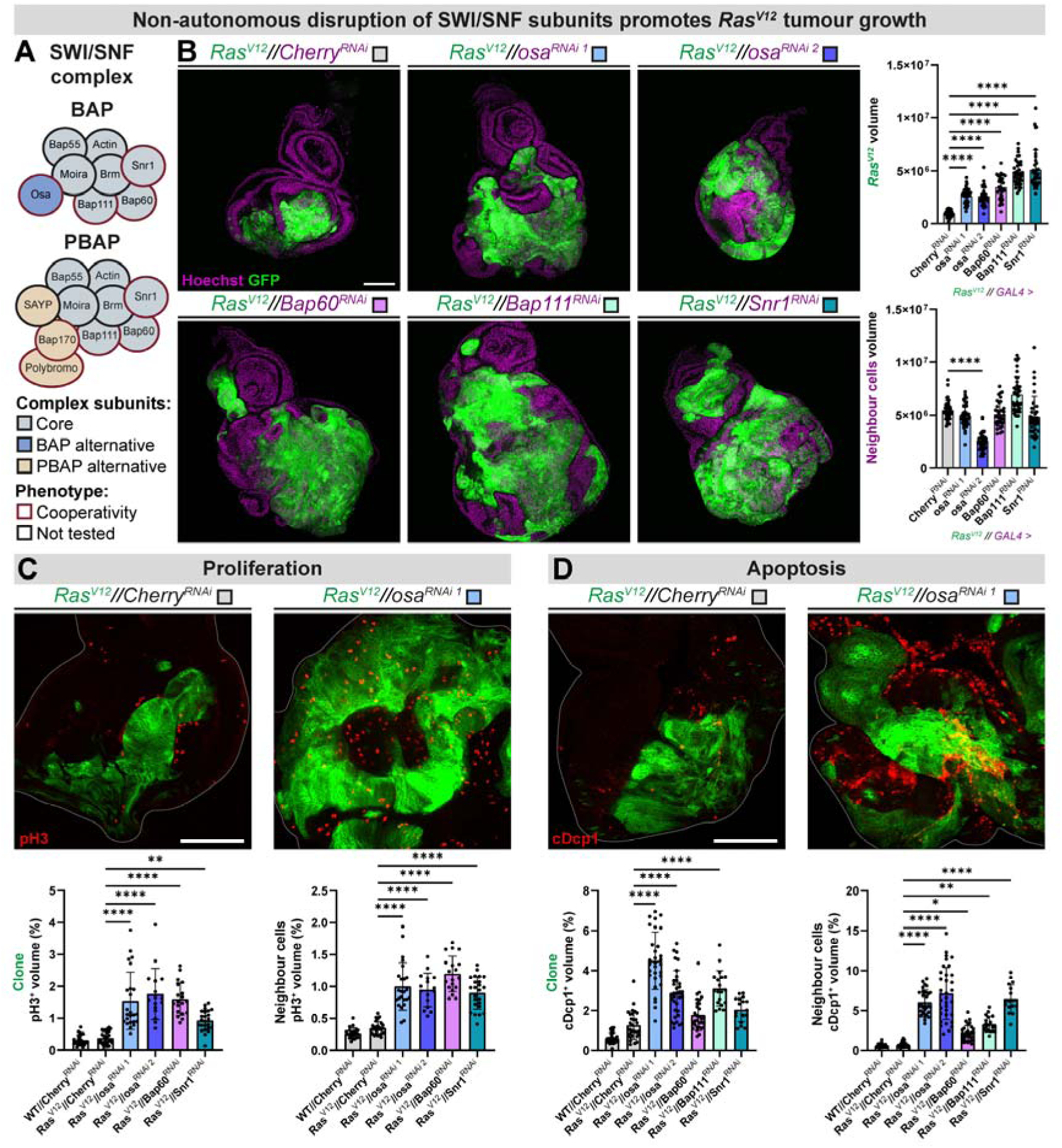
Disruption of SWI/SNF protein complex subunits in neighbouring cells enhances *Ras^V12^* clonal growth and proliferation, despite promoting apoptosis. (A) Schematic representation of the *Drosophila* SWI/SNF chromatin-remodelling complexes BAP and PBAP, and their core and alternative subunits. Subunits that generate an interclonal cooperative effect upon knockdown are encircled in red. (B) Representative confocal images of EADs showing control (*Ras^V12^//Cherry^RNAi^*, n=42) and downregulation of different SWI/SNF subunits in *Ras^V12^* neighbouring cells (*Ras^V12^//osa^RNAi1^*, n=41; *Ras^V12^//osa^RNAi2^*, n=39; *Ras^V12^//Bap60^RNAi^*, n=36; *Ras^V12^//Bap111^RNAi^*, n=39; and *Ras^V12^//Snr1^RNAi^*, n=37). In each condition, *Ras^V12^* cells are labelled with GFP (green), and nuclei are stained with Hoechst (magenta). Quantification of *Ras^V12^* clone volume (top right) and neighbouring tissue volume (bottom right) of distinct EADs is shown (µm^3^). Statistical significance was assessed with Kruskal–Wallis and Dunn’s test. Scale bar = 100 µm. (C) Representative confocal images of EADs and quantification of pH3-positive volume fraction within GFP^+^ clones and in the neighbouring cells of distinct EADs in the genotypes of WT control (*WT//Cherry^RNAi^*, n=23), *Ras^V12^* control (*Ras^V12^//Cherry^RNAi^*, n=27), and upon non-autonomous downregulation of SWI/SNF subunits (*Ras^V12^//osa^RNAi1^,* n=24; *Ras^V12^//osa^RNAi2^,* n=16; *Ras^V12^//Bap60^RNAi^,* n=19; and *Ras^V12^//Snr1^RNAi^,* n=24). In each condition, *Ras^V12^* cells are labelled with GFP (green), and EAD are stained for pH3 (red). Statistical significance was determined with Kruskal–Wallis and Dunn’s test. Scale bar = 100 µm. (D) Representative confocal images of EADs and quantification of cDcp1-positive volume fraction within GFP^+^ clones and in the neighbouring cells of distinct EADs in the genotypes of WT control (*WT//Cherry^RNAi^*, n=27), *Ras^V12^* control (*Ras^V12^//Cherry^RNAi^*, n=30), and upon non-autonomous downregulation of SWI/SNF subunits (*Ras^V12^//osa^RNAi1^,* n=30; *Ras^V12^//osa^RNAi2^,* n=34; *Ras^V12^//Bap60^RNAi^,* n=30; *Ras^V12^//Bap111^RNAi^,* n=19; and *Ras^V12^//Snr1^RNAi^,* n=17). In each condition, *Ras^V12^* cells are labelled with GFP (green), and EAD are stained for cDcp1 (red). Statistical significance was determined with Kruskal–Wallis and Dunn’s test. Scale bar = 100 µm.

Several studies have identified SWI/SNF variant-specific roles^51–55^. However, in the context of *Ras^V12^*clonal growth, we observe that the disruption of either BAP-specific (*Ras^V12^//osa^RNAi^)* or PBAP-specific (*Ras^V12^//Bap170^RNAi^, Ras^V12^//polybromo^RNAi^*) subunits in the surrounding cells promotes non-autonomous growth of *Ras^V12^* clones (Fig. 2A, 4B). Furthermore, downregulation of core SWI/SNF subunits (*Ras^V12^//Bap60^RNAi^, Ras^V12^//Bap111^RNAi^,* and *Ras^V12^//Snr1^RNAi^*) also triggered interclonal cooperativity (Fig. 4B). Among these, non-autonomous knockdown of core subunits Bap111 and Snr1 had the most pronounced effect on *Ras^V12^* tumour size (Fig. 4B). This data suggests that BAP and PBAP complexes may potentially partially compensate for one another for its tumour-suppressing activity.

Moreover, we did not observe significant changes in the volume of the SWI/SNF-deficient compartment, with the exception of one out of three independent *osa*^RNAi^ lines (Fig. 2A, 4B – *osa^RNAi^ ^2^*). Notably, the *osa^RNAi2^* line that reduced compartment size induced a non-autonomous increase in *Ras^V12^* tumour growth comparable to that observed with another *osa*^RNAi^ ^1^ line that did not affect compartment volume (Fig. 4B). This shows that the non-autonomous effect on *Ras^V12^* overgrowth is uncoupled from, and not a result of neighbouring compartment size reduction.

We further characterised the effects of SWI/SNF subunit loss on proliferation and apoptosis (Fig. 4C, D). Non-autonomous downregulation of SWI/SNF components Osa, Bap60, and Snr1 significantly increased proliferation, as measured through the mitotic marker pH3, both within the SWI/SNF-deficient neighbouring cells and *Ras^V12^* clones, when compared to control (Fig. 4C). The enhanced proliferation observed within *Ras^V12^* clones was consistent with the increased *Ras^V12^*clone volume (Fig. 4B, C).

Despite the increased proliferation, the overall volume of SWI/SNF-deficient tissue compartments remained unchanged, suggesting a balance between cell division and cell loss. Previous studies have shown that SWI/SNF loss in the wing imaginal disc leads to increased death^49,50^. Indeed, staining for cleaved Dcp1 caspase (cDcp1) revealed a strong increase in apoptosis in both SWI/SNF-deficient cells and *Ras^V12^* clones (Fig. 4D). Although apoptosis was elevated within *Ras^V12^* clones, this was not accompanied by a reduction or maintenance of clone size, but on the contrary, clone volume increased (Fig. 4B, D). This observation aligns with previous findings in *Ras^V12^, scrib^-^* tumours, where caspase activity has been shown to support tumour growth^56^. It is thus possible that this caspase activity can promote the death of SWI/SNF-deficient cells and simultaneously have a tumour growth-promoting function in neighbouring *Ras^V12^* clones.

We next asked whether the SWI/SNF tumour-promoting function act solely non-autonomously from neighbouring cells or can act from within *Ras^V12^*-transformed cells. Strikingly, depletion of SWI/SNF subunits in *Ras^V12^* tumours did not lead to changes in tumour growth, nor any systematic increase in proliferation (Suppl. fig. 3A, B). In contrast, SWI/SNF disruption, generally increased apoptosis in the *Ras^V12^* tumour compartment when knocked down within transformed cells (Suppl. fig. 3C). These results suggest that disruption of TSG SWI/SNF components carries higher tumourigenic potential when outsourced to neighbouring clones to promote tumour growth.

Together, these results show that disruption of either variant-specific or core subunits of the SWI/SNF complex in neighbouring cells promotes the non-autonomous growth of *Ras^V12^* clones. This behaviour is not supported when SWI/SNF depletion and *Ras^V12^*oncogene expression occur in the same compartment.

### Loss of SWI/SNF subunits in epithelial cells induces a wound-like inflammatory response

Non-autonomous disruption of the SWI/SNF subunit Osa induces a striking 2.7-fold increase in *Ras^V12^* tumour growth within five days (Fig. 4B). To uncover regulators of this interclonal cooperative behaviour we performed three bulk RNA-sequencing (RNAseq) experiments (Fig. 5A).

**Fig. 5.**
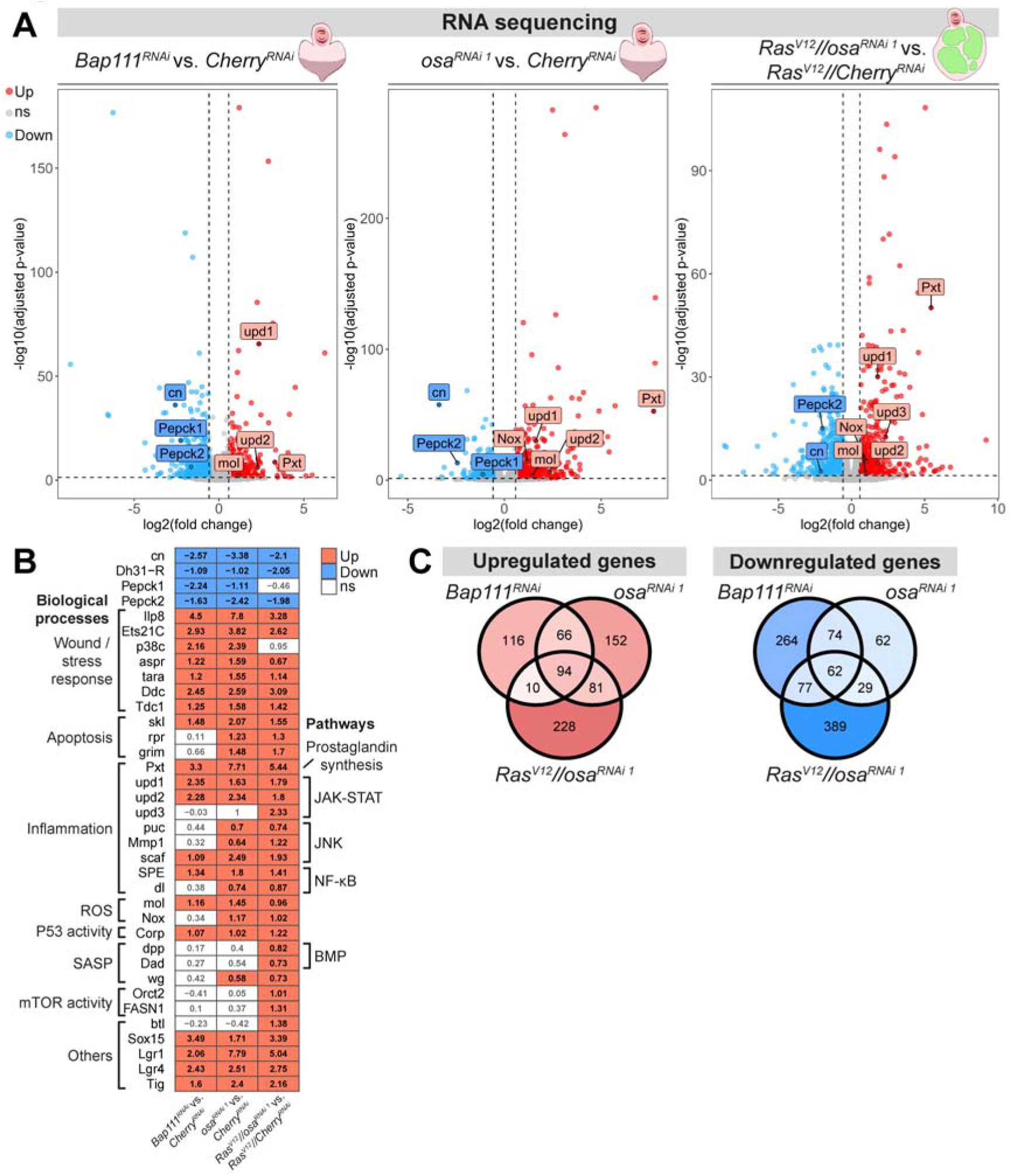
Transcriptome profiling identifies that disruption of SWI/SNF subunits in epithelia triggers a wound-like inflammatory stress response. (A) Volcano plots display genes found to be differentially expressed (|Log2FC| threshold ≥ 0.58 and an adjusted p-value threshold of ≤0.05) after bulk RNA sequencing of EAD in the following conditions: whole-EAD knockdown of *Bap111* (*Bap111^RNAi^* vs. *Cherry^RNAi^*), whole-EAD knockdown of *osa* (*osa^RNAi^ ^1^* vs. *Cherry^RNAi^*), or knockdown of *osa* in *Ras^V12^* neighbouring cells (*Ras^V12^//osa^RNAi^ ^1^* vs. *Ras^V12^//Cherry^RNAi^*), all compared with respective controls. Upregulated genes are labelled in red, downregulated genes are labelled in blue, and non-significantly changing genes are labelled in grey. Genes of interest found to be differentially expressed in one or more of the datasets are labelled in the volcano plot. (B) Heatmap shows differentially expressed status (Upregulated in red, downregulated in blue, and non-significant in white, |Log2FC| threshold ≥ 0.58 and an adjusted p-value threshold of ≤0.05) for genes of interest in the categories of Wound/stress response, apoptosis, inflammation, ROS generation, P53 activity, SASP, mTOR activity, and others, across the different datasets *Bap111^RNAi^* vs. *Cherry^RNAi^*, *osa^RNAi^ ^1^* vs. *Cherry^RNAi^*, and *Ras^V12^//osa^RNAi^ ^1^* vs. *Ras^V12^//Cherry^RNAi^.* (C)Venn diagrams show the intersection of upregulated and downregulated genes (|Log2FC| threshold ≥ 0.58 and an adjusted p-value threshold of ≤0.05) across the different datasets *Bap111^RNAi^* vs. *Cherry^RNAi^*, *osa^RNAi^ ^1^* vs. *Cherry^RNAi^*, and *Ras^V12^//osa^RNAi^ ^1^* vs. *Ras^V12^//Cherry^RNAi^.* Venn diagram compartments colour intensity follows a gradient based on the number of genes within the compartment.

In the first two experiments, we downregulated two different subunits of the SWI/SNF complex (*osa* and *Bap111*) throughout the entire EAD and compared their transcriptomes to those of *Cherry^RNAi^* control discs (Fig. 5A). With this approach, we aimed to identify genes that are commonly altered in expression upon disruption of the SWI/SNF complex, regardless of which subunit is targeted. Additionally, we performed bulk RNAseq of EADs containing *Ras^V12^* tumours surrounded by *osa^RNAi^* neighbour cells and compared them to EAD with *Ras^V12^* tumours alone (Fig. 5A). Our rationale was that genes differentially expressed only in the interclonal context (3^rd^ dataset), but not in EADs with uniform SWI/SNF disruption (1^st^ and 2^nd^ datasets), likely represent transcriptional changes specific to *Ras^V12^* tumours in response to SWI/SNF-deficient neighbouring cells, or vice versa.

By intersecting the differentially expressed genes across the three datasets, we found 66 genes commonly upregulated between *Bap111^RNAi^* and *osa^RNAi^* conditions, and 94 genes that were consistently upregulated across all three conditions, including *Ras^V12^//osa^RNAi^* (Fig. 5C). Conversely, we have identified 74 genes commonly downregulated between *Bap111^RNAi^* and *osa^RNAi^* samples, and 62 genes commonly downregulated in all three conditions (Fig. 5C). Among the consistently upregulated genes in the three datasets, we detected an increase in *upd1*, *upd2* / IL-6 (Fig. 5A, B), the ligands responsible for activating JAK-STAT signalling downstream of the Dome receptor (IL6 receptor-like). Indeed, JAK-STAT signalling has been previously shown to be increased upon SWI/SNF loss in the wing imaginal disc^49^, thus validating our transcriptomic approach. Curiously, the third gene encoding a JAK-STAT ligand, *upd3*, which has been described to mediate specific systemic effects^57–59^, was only found to be elevated in *Ras^V12^//osa^RNAi^* condition (Fig. 5A, B).

Transcriptome analysis revealed that several genes known to play a role during stress responses, such as wounding regeneration, were upregulated upon SWI/SNF disruption. Consistent with a wound-like scenario, and the increased apoptosis seen above (Fig. 4D), we observed the upregulation of pro-apoptotic genes *sickle* (skl)^60–62^ in all three conditions, and *reaper* (rpr)^63^ and *grim*^64^ in *osa^RNAi^*and *Ras^V12^//osa^RNAi^* samples (Fig. 5B). Furthermore, several genes known to be upregulated and/or to play a role during the wounding stress response were found to be upregulated, *Insulin-like peptide 8* (Ilp8)^65–69^, the transcription factor *Ets21c*^70^, *p38c* mitogen-activated protein kinase (MAPK)^71,72^, *asperous* (aspr)^70,73^, *taranis* (tara)^74^ melanization-promoting *Dopa decarboxylase* (Ddc)^75^, and *Tyrosine decarboxylase 1* (Tdc1)^76^ (Fig. 5B).

Beyond detecting inflammatory gene signatures associated with JAK-STAT pathway activation, we also observed increased expression of JNK signalling targets, such as *puckered* (puc)^77,78^, *Matrix metalloproteinase 1* (Mmp1)^79^, and *scarface* (scaf)^80,81^ (Fig. 5B). In addition, components of the NFκB pathway were upregulated, including *Spätzle-processing enzyme* (*SPE*) and *dorsal* (*dl*) (Fig. 5B). Notably, disruption of SWI/SNF also led to strong upregulation of *Peroxinectin-like* (Pxt) (Fig. 5B), a cyclooxygenase (COX)-like enzyme that has been described to act as mammalian *Cox1*, the rate-limiting enzyme in prostaglandin (PG) synthesis^82,83^. PGs are lipid signalling molecules that can be secreted to bind different receptors in an autocrine or paracrine manner. Although the role of PGs in *Drosophila* remains comparatively underexplored and has primarily been linked to development and female reproduction^84–86^, PGs are well established mediators of inflammation in mammals and are induced in response to tissue injury^87,88^.

Moreover, *Companion of reaper* (Corp), the Mdm2-like transcriptional target of p53^89^, was found to be upregulated upon SWI/SNF loss (Fig. 5B), suggesting there could be an ongoing DNA damage response. We detected a reactive oxygen species (ROS) production signature, evidenced by the upregulation of *NADPH oxidase* (Nox) and the Duox maturation factor *moladietz* (mol)^90,91^ (Fig. 5B).

A set of genes also showed a conserved downregulation response in the three datasets, such as *cinnabar* (cn)/kynurenine 3-monooxygenase, *Diuretic hormone 31 Receptor* (Dh31-R), and *Phosphoenolpyruvate carboxykinase 1* (Pepck1) and *Pepck2* (Fig. 5B).

Moreover, we identified several genes that were specifically upregulated in the *Ras^V12^//osa^RNAi^* condition, which may represent genes that are transcriptionally induced in *Ras^V12^* tumours in response to neighbouring SWI/SNF-deficient cells. Among these, we found increased expression of two known mTOR downstream targets required for growth, *caldéron* (Orct2)^92^ and *FASN1*^93,94^ (Fig. 5B). Additional genes in this category included components of the BMP pathway, *decapentaplegic* (*dpp*) and *Daughters against dpp* (*Dad*), as well as the FGF receptor *breathless* (*btl*) (Fig. 5B).

Together, these results suggest that disruption of SWI/SNF subunits triggers a complex wound-like inflammatory environment in the EAD. Given that wound regeneration relies on programs orchestrating collective cell behaviours through paracrine signals, this altered environment likely drives the non-autonomous overgrowth of adjacent *Ras^V12^* tumours.

### SWI/SNF loss in neighbouring cells promotes inflammation and an anabolic reprogramming of *Ras^V12^*clones

To determine whether the transcriptional changes identified by RNAseq are reflected *in vivo*, we next assessed inflammatory, stress, and anabolic pathway activity in the EAD following SWI/SNF disruption. Using established genetic reporters and immunostainings, we examined JAK-STAT and JNK signalling, ROS levels, DNA damage, and anabolic activity, as measured by mTOR activation and translation levels (Fig. 6).

**Fig. 6.**
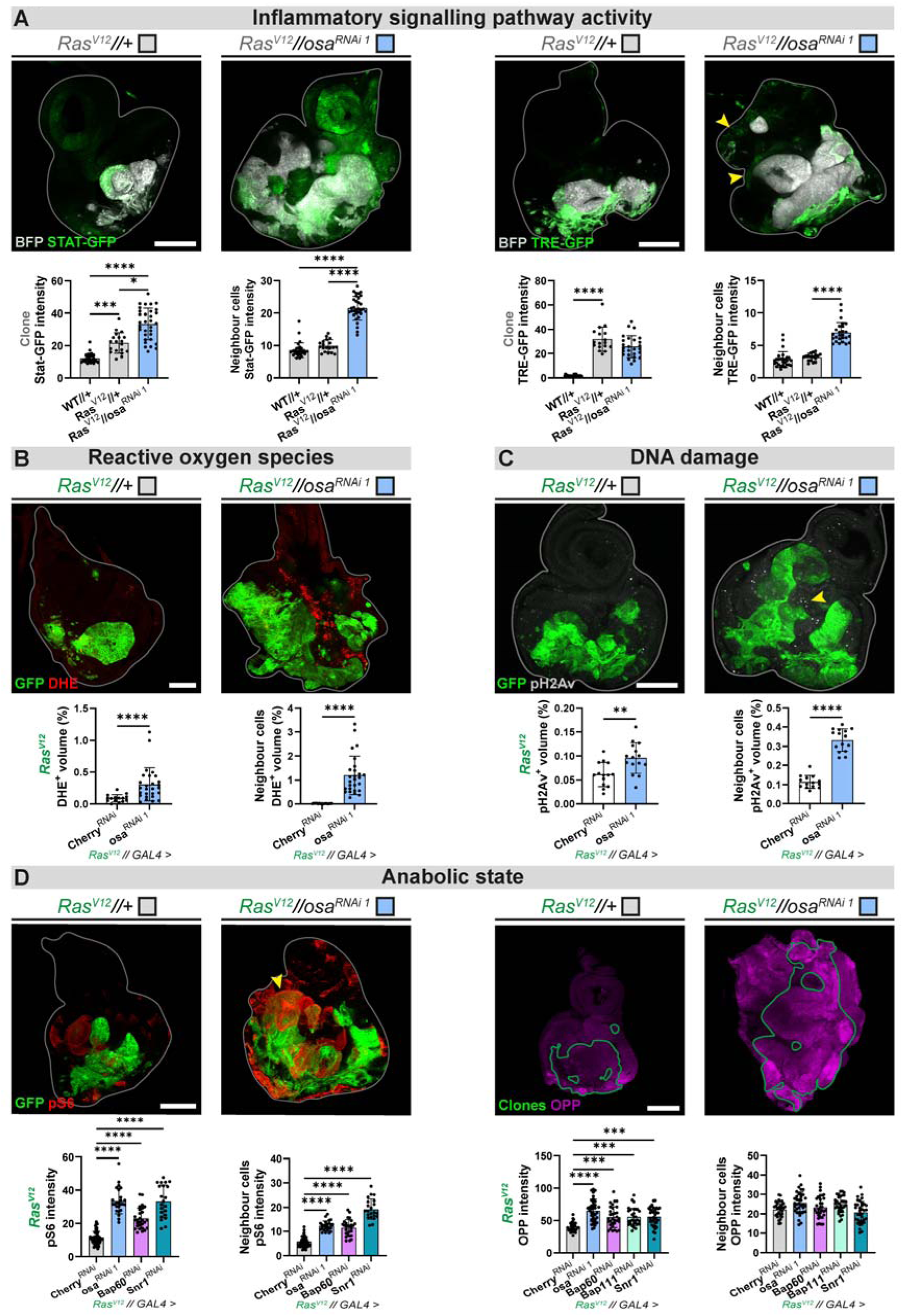
Non-autonomous SWI/SNF disruption promotes inflammatory signalling, cellular stress, and anabolic rewiring of *Ras^V12^* tumours. (A) Inflammatory signalling activity upon *osa* knockdown in *Ras^V12^* neighbouring cells. Representative confocal images of EADs and quantification of STAT-GFP and TRE-GFP fluorescent intensity (mean grey value) within GFP^+^ clones and in the neighbouring cells of distinct EADs in the genotypes of WT control (*WT//+*, STAT-GFP – n=31, TRE-GFP – n=28), *Ras^V12^* control (*Ras^V12^//+*, STAT-GFP – n=19, TRE-GFP – n=17), and upon non-autonomous downregulation of osa (*Ras^V12^//osa^RNAi1^*, STAT-GFP – n=32, TRE-GFP – n=26). In each condition, *Ras^V12^* cells are labelled with BFP (grey), and enhancer reporter activity is shown with GFP (green). Statistical significance between groups was determined with Kruskal–Wallis and Dunn’s test. Scale bar = 100 µm. (B) Intracellular ROS levels upon *osa* knockdown in *Ras^V12^* neighbouring cells. Representative confocal images of EADs and quantification of DHE^+^ volume fraction (%) within *Ras^V12^* clones and in the neighbouring cells of distinct EADs in the genotypes of *Ras^V12^* control (*Ras^V12^//Cherry^RNAi^*, n=15), and upon non-autonomous downregulation of osa (*Ras^V12^//osa^RNAi^ ^1^*, n=27). In each condition, *Ras^V12^* cells are labelled with GFP (green), and DHE (red). Statistical significance between groups was determined with Mann–Whitney U test. Scale bar = 100 µm. (C) DNA damage response upon *osa* knockdown in *Ras^V12^* neighbouring cells. Representative confocal images of EADs and quantification of pH2Av^+^ volume fraction (%) within *Ras^V12^* clones and in the neighbouring cells of distinct EADs in the genotypes of *Ras^V12^* control (*Ras^V12^//Cherry^RNAi^*, n=14), and upon non-autonomous downregulation of *osa* (*Ras^V12^//osa^RNAi1^*, n=15). In each condition, *Ras^V12^* cells are labelled with GFP (green), and pH2Av (grey). Statistical significance between groups was determined with Welch’s t-test. Scale bar = 100 µm. (D) Tissue anabolic rewiring upon *osa* knockdown in *Ras^V12^* neighbouring cells. Representative confocal images of EADs and quantification of pS6 levels (mean grey value) within *Ras^V12^* clones and in the neighbouring cells of distinct EADs in the genotypes of *Ras^V12^* control (*Ras^V12^//Cherry^RNAi^*, n=49), and upon non-autonomous downregulation of SWI/SNF subunits (*Ras^V12^//osa^RNAi1^*, n=25; *Ras^V12^//Bap60^RNAi^*, n=28; *Ras^V12^//Snr1^RNAi^*, n=22). Representative confocal images of EADs and quantification of OPP incorporation (mean grey value) within *Ras^V12^* clones and in the neighbouring cells of distinct EADs in the genotypes of *Ras^V12^* control (*Ras^V12^//Cherry^RNAi^*, n=29), and upon non-autonomous downregulation of SWI/SNF subunits (*Ras^V12^//osa^RNAi1^*, n=36; *Ras^V12^//Bap60^RNAi^*, n=32; *Ras^V12^//Bap111^RNAi^*, n=31; *Ras^V12^//Snr1^RNAi^*, n=35). For the pS6 experiment, *Ras^V12^* cells are labelled with GFP (green), and EAD are stained for pS6 (red). Quantification Statistical significance between groups was determined with Kruskal–Wallis and Dunn’s test. For the OPP experiment, *Ras^V12^* GFP-labelled cells are outlined (green), and OPP incorporation is shown (magenta). Statistical significance between groups was found with one-way ANOVA and Tukey’s test. Scale bar = 100 µm.

Consistent with a wound-like inflammatory state induced by SWI/SNF disruption, we observed elevated inflammatory signalling activity. Indeed, JAK-STAT signalling was elevated in Osa-depleted cells, as well as in neighbouring *Ras^V12^* clones compared with controls (Fig. 6A). JNK signalling activity revealed a consistent pattern in control samples, where mostly the posterior region of the *Ras^V12^*tumour exhibited elevated JNK signalling, while the remainder of the tumour showed minimal activity (Fig. 6A). A similar compartmentalized distribution was also observed for JAK-STAT signalling, although in distinct regions (Fig. 6A). This spatial segregation is consistent with findings by Jaiswal *et al.* that showed that during both regeneration and tumourigenesis, JNK and JAK-STAT pathways mutually repress each other, enabling the segregation of senescent and proliferative cell populations^95^. Upon downregulation of Osa in the surrounding tissue, we observed increased JNK activity only in the SWI/SNF-deficient cells, with no change in JNK activity in *Ras^V12^* tumour cells (Fig. 6A). These results support a model similar to that described in *Ras^V12^*//*scrib*^-^ context, where JNK activation in *scrib^-^* cells induces the expression of Upd/IL-6 cytokines, which in turn activates JAK-STAT signalling in neighbouring *Ras^V12^* tumour cells through a paracrine mechanism^26^.

Given the upregulation of the genes involved in ROS production, *mol* and *Nox*, and the p53 target *Corp*, in our RNAseq data (Fig. 5B), we next asked whether the inflammatory signalling induced by SWI/SNF disruption is associated with increased cellular stress, such as oxidative stress and DNA damage (Fig. 6B, C). Indeed, using Dihydroethidium (DHE) staining to detect intracellular ROS, we observed a pronounced ROS increase in the Osa-depleted compartment compared with control (Fig. 6B). Although more modest, ROS levels were also elevated within *Ras^V12^* clones when surrounded by SWI/SNF-disrupted neighbouring cells (Fig. 6B). Moreover, staining for phosphorylated Histone H2Av (pH2Av), a marker of DNA double-strand breaks, revealed a similar spatial pattern of increased DNA damage in both the SWI/SNF and *Ras^V12^* clone compartments when compared to control (Fig. 6C).

Furthermore, the strong increase in *Ras^V12^* growth observed in the presence of neighbouring SWI/SNF-deficient cells suggests *Ras^V12^* tumours adopt a highly anabolic state to sustain their expansion. The mTOR complex is a central regulator of cellular metabolism that integrates external cues, such as growth factors, nutrient availability and energy status. When activated, mTOR promotes several anabolic processes, including protein, nucleotide, and lipid synthesis, while simultaneously suppressing catabolism through inhibition of autophagy^96^. Notably, mTOR is required for *RAS*-driven tumourigenesis in *in vivo* mouse models, and its hyperactivation further enhances oncogenic RAS tumour progression^97,98^. A key downstream effector of mTOR is the S6 kinase (S6K), which phosphorylates ribosomal protein S6. To assess tissue spatial mTOR activity, we used an antibody against phosphorylated S6 (pS6), a well-established readout of mTOR signalling, and which has been validated in *Drosophila* imaginal discs^99^. When we stained *Ras^V12^* only EAD, we observed a non-autonomous pS6 pattern that did not match the previously reported staining of WT EAD^100^. In *Ras^V12^* only EAD, several wildtype neighbour cells encircling the tumours showed elevation of pS6 (Fig. 6D), suggesting there is a mTOR microenvironmental response upon the presence of *Ras^V12^* tumours. Strikingly, non-autonomous downregulation of SWI/SNF subunits Osa, Bap60, and Snr1, resulted in the increase of pS6 intensity in both the surrounding SWI/SNF-deficient tissue and inside *Ras^V12^* tumours (Fig. 6D). Notably, pS6 staining within the *Ras^V12^* tumours was not uniform, but rather showed compartmentalized activation (Fig. 6D). Recently, it has been described that JAK-STAT signalling can bypass insulin signalling to promote anabolism for regeneration upon wounding^101^. Given that JAK-STAT activity is also spatially compartmentalized, it is plausible that regional JAK-STAT signalling mediates mTOR activation in discrete subpopulations within *Ras^V12^* clones.

To confirm that the elevated mTOR activity was associated with increased anabolic behaviour, we performed an O-propargyl-puromycin (OPP) incorporation assay to measure translational activity. Strikingly, upon non-autonomous downregulation of SWI/SNF subunits, we observed a robust increase in protein synthesis within *Ras^V12^* clones, while translation levels in the surrounding SWI/SNF-deficient cells remained unchanged (Fig. 6D).

These results show that non-autonomous loss of SWI/SNF induces inflammatory signalling and anabolic reprogramming in *Ras^V12^* tumours, which likely supports tumour growth.

### Non-autonomous induction of *Ras^V12^* overgrowth requires ROS and prostaglandin synthesis machinery in SWI/SNF-deficient surrounding cells

Our transcriptomic and *in vivo* analyses implicated ROS production and prostaglandin signalling as key components of the inflammatory microenvironment induced by SWI/SNF disruption. We therefore tested whether these pathways are required within SWI/SNF-deficient neighbouring cells to drive non-autonomous *Ras^V12^* tumour overgrowth.

ROS signalling is widely employed by stressed cells to activate signalling pathway programs, such as the JNK pathway^72,91,102^, which could potentially trigger the release of pro-tumourigenic paracrine factors. Enzymatic detoxification of intracellular ROS through overexpression of cytosolic *Superoxide dismutase 1* (Sod1) in SWI/SNF disrupted cells limited the growth of neighbouring *Ras^V12^* tumours (Fig. 7A, B). This also led to a reduction in the size of SWI/SNF-deficient compartment (Fig. 7B).

**Fig. 7.**
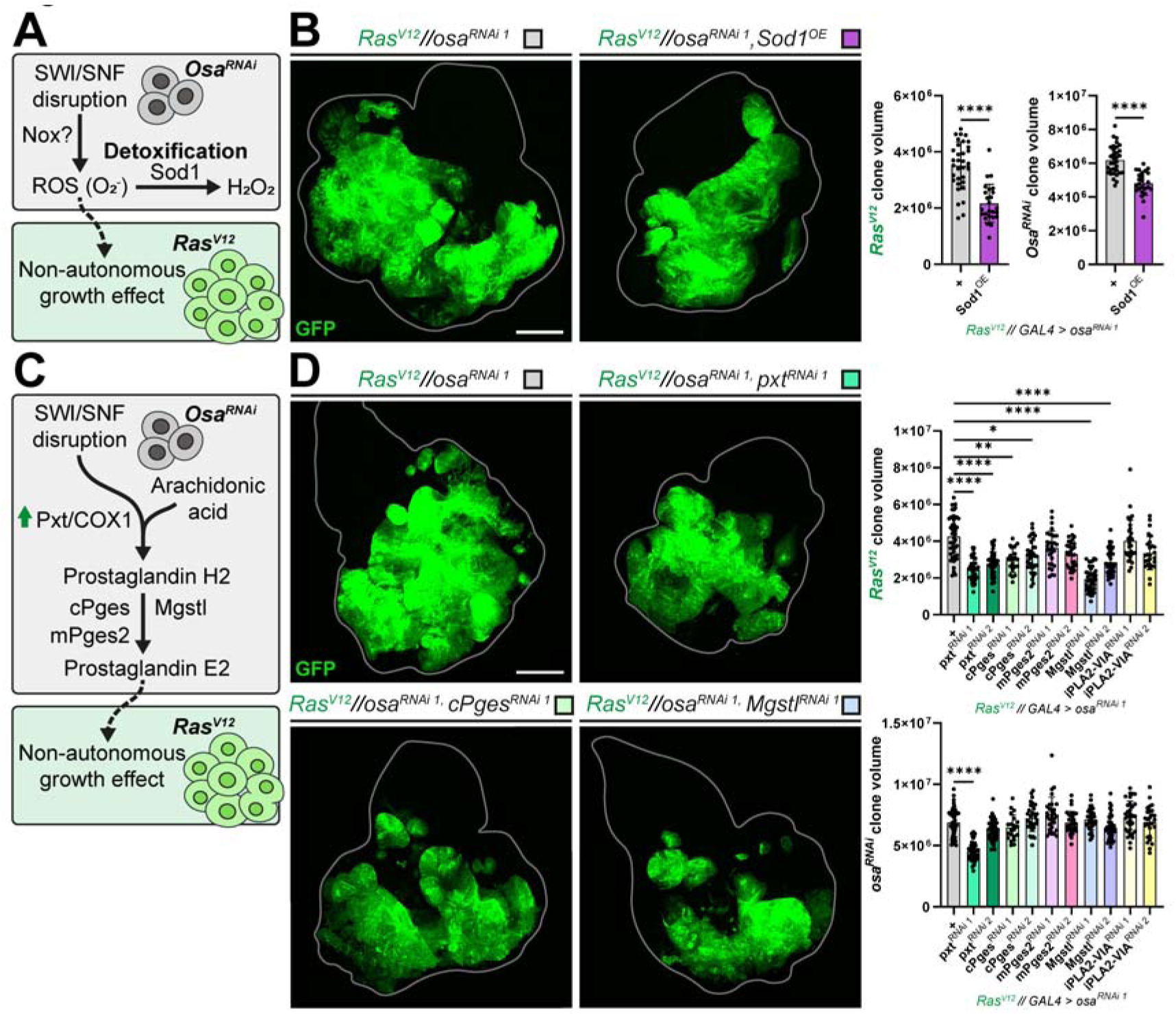
Prostaglandin synthesis machinery and ROS are required in SWI/SNF-deficient cells to interclonally promote *Ras^V12^* overgrowth. (A) Schematic illustrating a model in which disruption of SWI/SNF induces ROS production, leading to a non-autonomous growth effect on *Ras^V12^* tumours. Detoxification of superoxide can be achieved through Sod1. (B) Representative confocal images of EADs showing control (*Ras^V12^//osa^RNAi1^*, n=33) and overexpression of *Sod1* in *osa^RNAi^* cells neighbouring *Ras^V12^* tumours (*Ras^V12^//osa^RNAi1^, Sod1^OE^*, n=25). *Ras^V12^* cells are labelled with GFP (green), and disc outline is shown (grey). Quantification of *Ras^V12^* clone volume and neighbouring tissue volume of distinct EADs is shown (µm^3^). Statistical significance was assessed with Welch’s t-test. Scale bar = 100 µm. (C) Schematic illustrating a model in which disruption of SWI/SNF induces prostaglandin synthesis through upregulation of Pxt/Cox1, leading to a non-autonomous growth effect on *Ras^V12^* tumours. (D) Representative confocal images of EADs showing control (*Ras^V12^//osa^RNAi1^*, n=47) and downregulation of enzymes involved in prostaglandin synthesis in *osa^RNAi^* cells neighbouring *Ras^V12^* tumours (*Ras^V12^//osa^RNAi1^, pxt^RNAi1^*, n=39; *Ras^V12^//osa^RNAi1^, pxt^RNAi2^*, n=45; *Ras^V12^//osa^RNAi1^, cPges^RNAi1^*, n=21; *Ras^V12^//osa^RNAi1^, cPges^RNAi2^*, n=34; *Ras^V12^//osa^RNAi1^, mPges2^RNAi1^*, n=30; *Ras^V12^//osa^RNAi1^, mPges2^RNAi2^*, n=30; *Ras^V12^//osa^RNAi1^, Mgstl^RNAi1^*, n=41; *Ras^V12^//osa^RNAi1^, Mgstl^RNAi2^*, n=43; *Ras^V12^//osa^RNAi1^, iPLA2-VIA^RNAi1^*, n=32; *Ras^V12^//osa^RNAi1^, iPLA2-VIA^RNAi2^*, n=27). *Ras^V12^* cells are labelled with GFP (green), and disc outline is shown (grey). Quantification of *Ras^V12^* clone volume and neighbouring tissue volume of distinct EADs is shown (µm^3^). Statistical significance was assessed with Kruskal–Wallis and Dunn’s test. Scale bar = 100 µm.

A key rate-limiting step in prostaglandin synthesis is catalysed by COX enzymes, which convert arachidonic acid into prostaglandin H2 (PGH2). PGH2 is subsequently processed by downstream prostaglandin synthases to generate bioactive prostaglandins, such as prostaglandin E2 (PGE2), which mediate downstream signalling effects (Fig. 7C). We had previously observed that upon *osa* depletion, the *Drosophila* COX1-like enzyme *Pxt* was strongly upregulated (7.7 log2FC) compared to control EADs (Fig. 5B), suggesting that prostaglandin signalling may contribute to the interclonal cooperative behaviour. Strikingly, downregulation of *Pxt* in *osa^RNAi^* with two independent RNAis significantly restricted the non-autonomously induced *Ras^V12^*tumour overgrowth (Fig. 7D). Moreover, depletion in *osa^RNAi^* cells for two out three PGE2 synthase paralogues, *cytosolic Prostaglandin E synthase* (cPges) and *Microsomal glutathione S-transferase-like* (Mgstl), but not *microsomal Prostaglandin E synthase 2* (mPges2), resulted in a reduction of *Ras^V12^* overgrowth. Previous work in mammals has shown that the calcium-independent phospholipase A2, group 6 (PLA2G6) can be activated by effector caspase-3 and to source arachidonic acid (AA) for prostaglandin synthesis which stimulates tumour repopulation upon irradiation^103–105^. *Drosophila* encodes a homolog of *PLA2G6*, *iPLA2-VIA*, however, we did not observe any effects on *Ras^V12^* growth upon depletion of this lipase in *osa^RNAi^* cells (Fig. 7D). Recent work has shown that AA could potentially be sourced via *brummer*/*ATGL*^106^, suggesting that alternative lipid mobilisation pathways may supply AA for PG synthesis in SWI/SNF-deficient cells.

Together, these results demonstrate that ROS production and prostaglandin synthesis within SWI/SNF-deficient neighbouring cells are required to interclonally promote *Ras^V12^*tumour overgrowth.

## Discussion

Mutational mosaicism is a common phenomenon in both healthy^107^ and tumour tissues^108^, which arises from the accumulation of somatic mutations over time. This is particularly evident for Onc and TSGs, which often confer a selective fitness advantage, allowing the mutated clones to persist and expand within the tissue. While this genetic heterogeneity is a hallmark of aging organs and cancer evolution, our understanding of the functional consequences of this phenomenon are very limited. Specifically, the mechanisms by which interclonal interactions alter the relative abundance and behaviour of other neighbouring cell populations. This is particularly relevant in early tumourigenesis, where cooperative or competitive interactions could determine whether neighbouring tumour cells, regress, stay dormant, or progress into a full-blown tumour.

Although yet to be demonstrated in a clinical setting, it is likely that the fate of early *RAS*-mutant clones in humans is shaped by interclonal interactions. Despite being one of the most common mutational events in carcinomas, *RAS* mutations alone are often insufficient to drive cancer development and can even lead to the elimination of the mutant clone. Extrinsic phenomena, including interclonal interactions, have been shown in both *Drosophila* and mouse models to be strong determinants of *RAS*-driven tumourigenesis^26–28^. Yet, the interclonal interaction outcomes between *RAS* and other commonly mutated genes in human cancers had not been addressed. In this study, we performed a genetic screen to identify clinically relevant Onc and TSGs that can interclonally alter the growth behaviour of *Ras^V12^* tumours. To do this, we employed the *Drosophila* EyaHOST genetic system^34^, which enabled us to model interclonal interactions within an epithelial tissue *in vivo* and systematically assess the non-autonomous effects of misexpressing Onc and TSGs of interest. To the best of our knowledge, no other system offers a comparable combination of physiological relevance and throughput for modelling interclonal interactions.

Several genes identified to interclonally promote *Ras^V12^* tumour overgrowth encode components of the SWI/SNF chromatin remodelling complex. Disruption of either core or variant SWI/SNF subunits in neighbouring cells enhanced tumour expansion non-autonomously. As a consequence of being neighboured by SWI/SNF disrupted cells, *Ras^V12^* tumours proliferate more and undergo an anabolic shift, as observed through elevation of mTOR activity and protein translation. We observed that disruption of the SWI/SNF complex installs a wound-like microenvironment, characterised by increased caspase activity, elevation of inflammatory pathway activity (JAK-STAT, JNK, and PGs), and cellular stress (ROS and DNA damage). Notably, ROS and prostaglandin synthesis machinery were required in SWI/SNF disrupted cells to non-autonomously promote *Ras^V12^* tumour growth. Indeed, prostaglandins have been widely implicated in driving the progression of *RAS*-driven tumours^109–114^.

A recent study in mice showed that tissue injury induces pS6 in wound-adjacent tissue, seemingly through the release of damage-associated molecular patterns (DAMPs) release, and that this response is ultimately required for proper healing^115^. It is likely that disruption of SWI/SNF TSGs triggers a similar wound-like response, however, unlike transiently damaged tissue, these mutant clones are not cleared, potentially leading to a sustained and chronic activation of this program. Interestingly, a previous study in mice has shown that *osa*/*ARID1A* deletion leads to better regeneration of the liver and ear upon damage^116^. These findings suggest that SWI/SNF loss may prime tissues for a regenerating or stress responsive state, but which potentially could be exploited by neighbouring oncogenic clones to support tumour growth.

One question that remains unanswered in this study is why the loss of the SWI/SNF chromatin remodeller leads to a wound-like stress response. Besides remodelling chromatin to modulate transcription, SWI/SNF is also required for DNA replication and repair^117,118^. A possibility could be that failure to resolve these DNA processes could create an inflammatory response that could non-autonomously stimulate *Ras^V12^*growth. A recent study has shown in imaginal discs that replication stress causes DNA damage, activation of p53, inflammatory JNK activity, and ultimately apoptosis^119^. Another possibility is that SWI/SNF regulates damage-responsive enhancer regions, which control genes involved in wound repair^120,121^.

Moreover, a previous study has identified an interclonal suppressive behaviour for clones with deficient Notch signalling^33^. Here, we expand on this concept by uncovering several other additional Onc and TSGs that when disrupted can non-autonomously suppress *Ras^V12^* growth. Strikingly, several of these genes, such as *myc*, *tai*, and *ago*, have been identified to drive the elimination of nearby wildtype cells, a process termed supercompetition^42,43,47,48^. Our findings therefore reveal an unexpected vulnerability of *Ras^V12^* tumours to neighbouring supercompetitior clones, suggesting that altering the fitness of clones in the microenvironment shapes tumour progression.

Two of the genes identified in our screen to promote interclonal suppression, *vap* and *CycE*, have not previously been implicated in supercompetition. In the case of *vap,* which encodes the p120 Ras GAP, while seemingly counter-intuitive that a Ras overactivated neighbouring population via *vap^RNAi^* could be fitter than oncogenic *Ras^V12^* clones, this fitness imbalance may be explained by the different levels of Ras activation observed between the two insults. Lower levels of Ras activation have been proposed to be optimal for increased proliferation, while higher levels of Ras have been described to promote senescence^11,122–124^. Given that *vap^RNAi^* is expected to induce lower levels of MAPK signalling than *Ras^V12^*overexpression, it is possible that *vap^RNAi^* are more competitive than *Ras^V12^* cells by benefiting from MAPK activation without triggering tumour suppressive programs such as oncogene-induced senescence.

As for *CycE^OE^*, one possible mechanism by which these clones suppress *Ras^V12^* tumour growth may relate to their accelerated G1-to-S phase entry. Since *Ras^V12^* clones already exhibit elevated *CycE* levels and are prone to replication stress^125–127^, the increased nucleotide demand and uptake by neighbouring *CycE^OE^* cells could deplete the local extracellular nucleotide pool, thereby exacerbating replication stress in *Ras^V12^* clones and impairing growth. Supporting this idea, a recent study in imaginal discs showed that cells undergoing replication stress can be rescued by nucleotide sharing from their neighbouring cells via gap junctions^128^.

Although our study uncovers previously unrecognised interclonal interactions between clinically relevant TSGs/Onc and early *Ras^V12^* tumours, it does not address whether early human lesions present conserved clonal architectures that enable the interactions found here. Nevertheless, accumulating evidence from patient-derived data suggests that these interclonal behaviours could occur in human tumours. Immunohistochemical analyses indicate that heterogeneous loss of *ARID1A* (*osa* in *D. melanogaster*) is a recurrent feature across multiple tumour types, including endometrial^129,130^, ovarian^131,132^, non-small cell lung^133^, gastric^134,135^, and colorectal^136^ cancers. In colorectal cancer, heterogeneous *ARID1A* loss is particularly prevalent, with nearly 60% of *ARID1A*-mutant cases displaying heterogeneous loss rather than complete inactivation^136^. Notably, in gastric cancer, cases with partial *ARID1A* loss are associated with poorer disease-free and overall survival compared with those exhibiting uniform loss^134^. Together, these observations suggest that heterogeneous *ARID1A* loss is not only widespread but may actively contribute to tumour progression, potentially through interclonal interactions.

Furthermore, a key result in this study was that autonomous deficiency for SWI/SNF did not promote *Ras^V12^*growth, in contrast with the increased *Ras^V12^* clone expansion when this complex was disrupted in the surrounding cells. While *ARID1A* is commonly defined as a TSG and its inactivation is frequently associated to cancer progression, evidence indicates that its role is context-dependent, and that *ARID1A* loss does not universally correlate with adverse clinical outcomes^137^. In colorectal cancer, *ARID1A* loss has paradoxically been associated with a better prognosis^138^, or shown to have no impact on patient survival^139–142^. Although the mechanisms underlying these observations remain poorly understood, it could potentially be that *ARID1A* deficiency fails to cooperate autonomously with MAPK pathway mutations, which represent an important transformation event in colorectal tumours. Indeed, previous studies suggest that the combined disruption of SWI/SNF subunits and MAPK pathway activation does not induce autonomous cooperativity, and could constrain tumour growth. Sen *et al.* show that colorectal cancer cell lines with *ARID1A* loss and *KRAS^G13D^*have reduced proliferation in comparison to KRAS wildtype lines, and that *ARID1A* is required for MAPK-dependent transcription by enabling AP-1 enhancer activity^143^. Similarly, Malik *et al.* described that *SMARCA4*, the ortholog of *Drosophila brm*, is required for *KRAS^G12D^*-driven tumourigenesis in the mouse lung and human cells^144^. In line with this context dependency, Song *et al.* described in *Drosophila* that although autonomous disruption of SWI/SNF together with *yorkie* overexpression induces cooperative growth, it fails to enhance tumour development when combined with *Egfr* overexpression^50^. Collectively, these studies argue that disruption of SWI/SNF is incompatible with MAPK-driven tumourigenesis in an autonomous setting. Instead, our findings indicate that SWI/SNF disruption can promote MAPK-driven tumours if disrupted in the neighbouring cells. To our knowledge, this is the first report of a cancer driver whose tumour-promoting effects need to be outsourced to neighbouring cells, likely due to cell-intrinsic incompatibilities with the already existing oncogenic lesions. It is tempting to speculate that the inconsistent clinical outcomes observed in *ARID1A* mutant colorectal cancer patients could perhaps be caused by instances where the tumour contains *ARID1A* mutated autonomously versus non-autonomously.

In conclusion, our findings demonstrate that the clonal landscape is a critical determinant for tumourigenesis, with independent clones capable of promoting or suppressing the growth of neighbouring tumours. We expand the current understanding of the mutational drivers and mechanisms that govern interclonal interactions, which opens for future studies aiming to exploit interclonal crosstalk as potential therapeutic strategy in preclinical models.

**Suppl. fig. 1.**
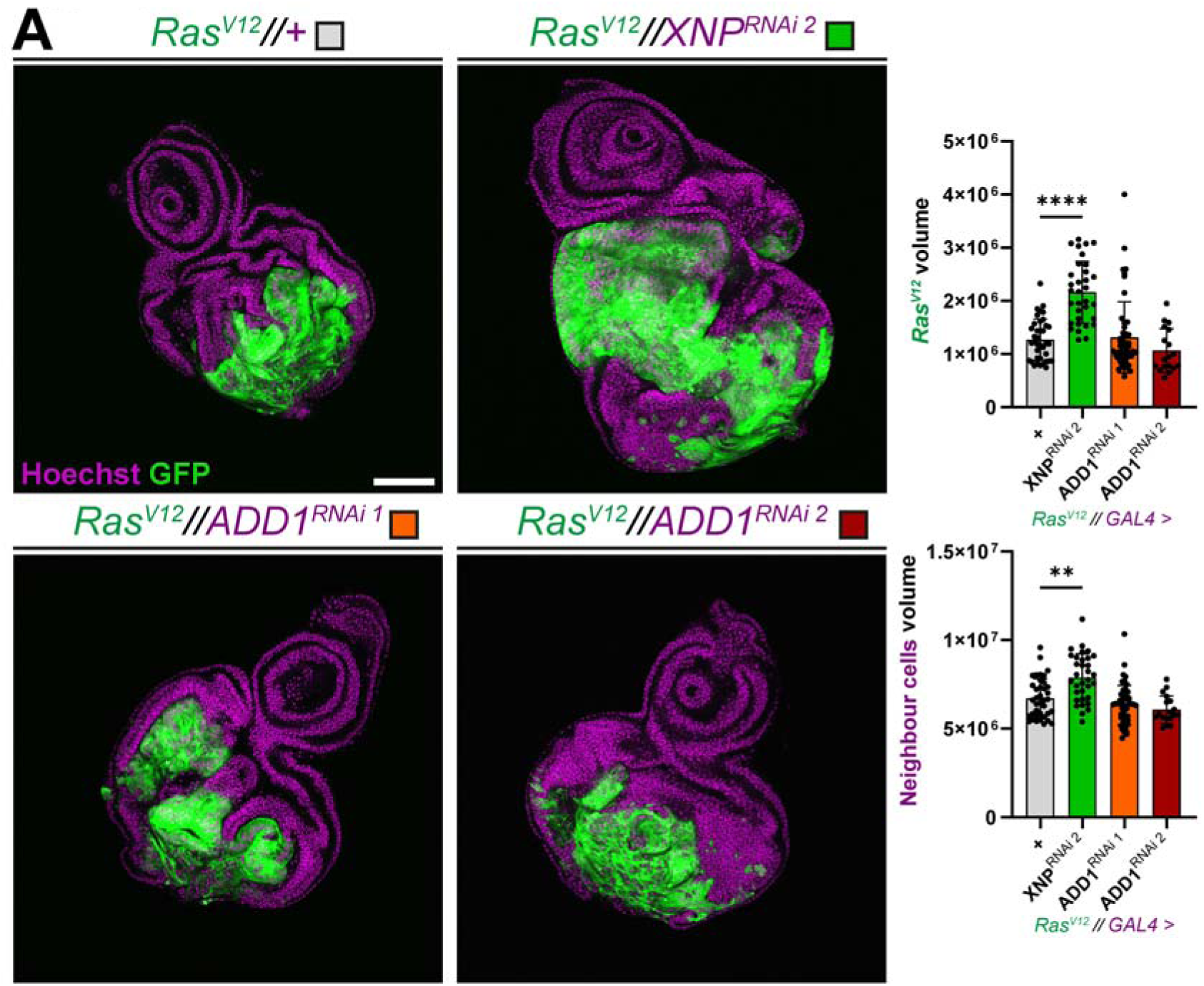
Loss of *XNP* in neighbouring cells, but not *ADD1*, non-autonomously promotes the growth of Ras^V12^ tumours. (A) Representative confocal images of EADs showing control (*Ras^V12^//+*, n=38) and the effect of knocking down in neighbouring cells *XNP* (*Ras^V12^//XNP^RNAi^ ^2^*, n=33) and *ADD1* (*Ras^V12^//ADD1^RNAi1^*, n=50*; Ras^V12^//ADD1^RNAi2^*, n=18). In each condition, *Ras^V12^* cells are labelled with GFP (green), and nuclei are stained with Hoechst (magenta). Quantification of *Ras^V12^*clone volume (top right) and neighbouring tissue volume (bottom right) of distinct EADs is shown (µm^3^). Statistical significance between groups was tested with Kruskal–Wallis and Dunn’s test. Scale bar = 100 µm.

**Suppl. fig. 2.**
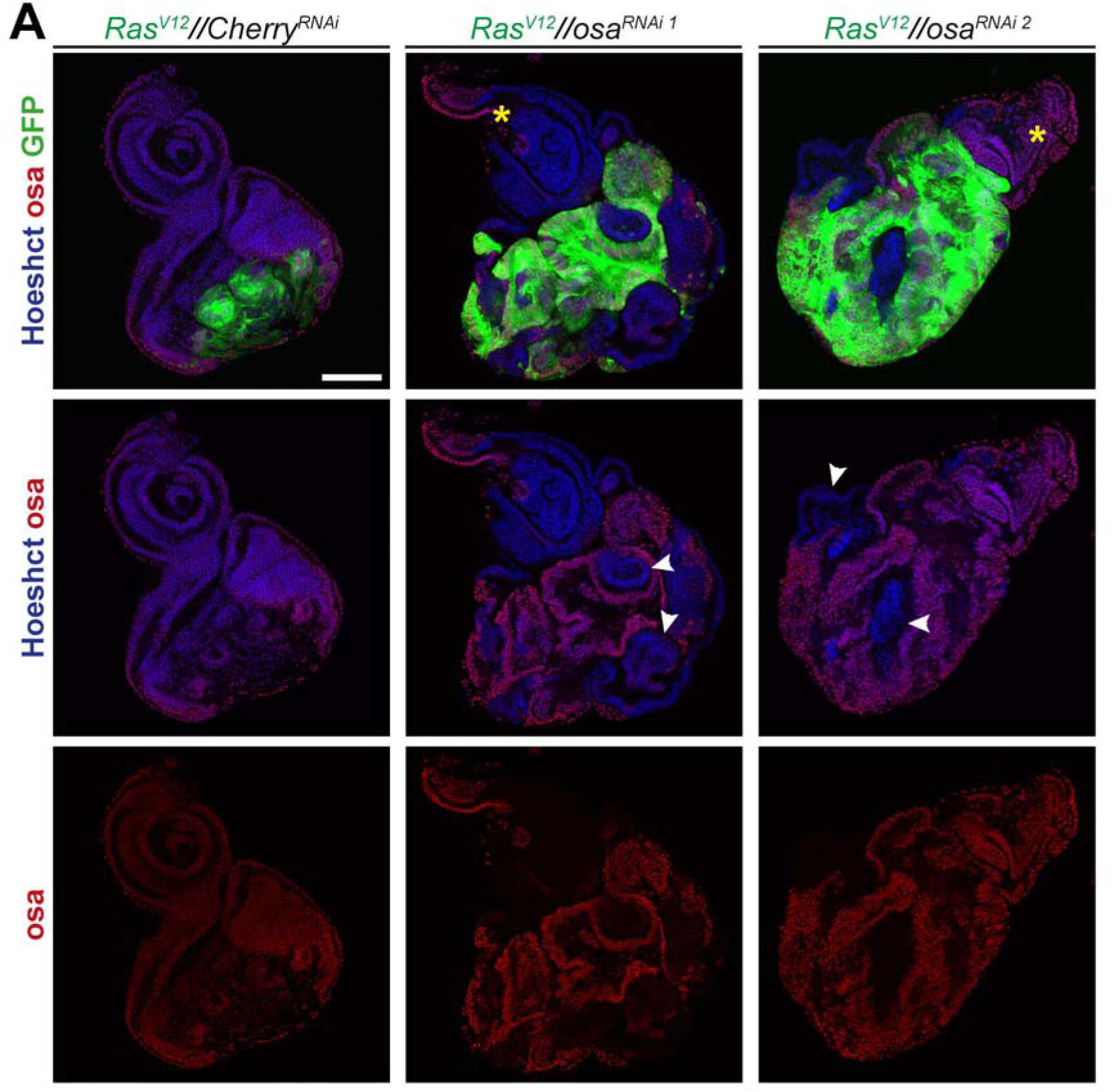
Validation of osa knockdown effect in *Ras^V12^* tumour surrounding cells. (A) Representative confocal images of EADs showing control (*Ras^V12^//Cherry^RNAi^*) and downregulation of *osa* in *Ras^V12^* neighbouring cells (*Ras^V12^//osa^RNAi1^, Ras^V12^//osa^RNAi2^*). In each condition, *Ras^V12^* cells are labelled with GFP (green), stained for osa (red), and nuclei are stained with Hoechst (blue). White arrows indicate *Ras^V12^*neighbouring cells where downregulation of osa protein is evident. Yellow asterisk indicates regions in the antenna portion of the disk, which are commonly found unflipped by ey-flp with the EyaHOST system. Scale bar = 100 µm.

**Suppl. fig. 3.**
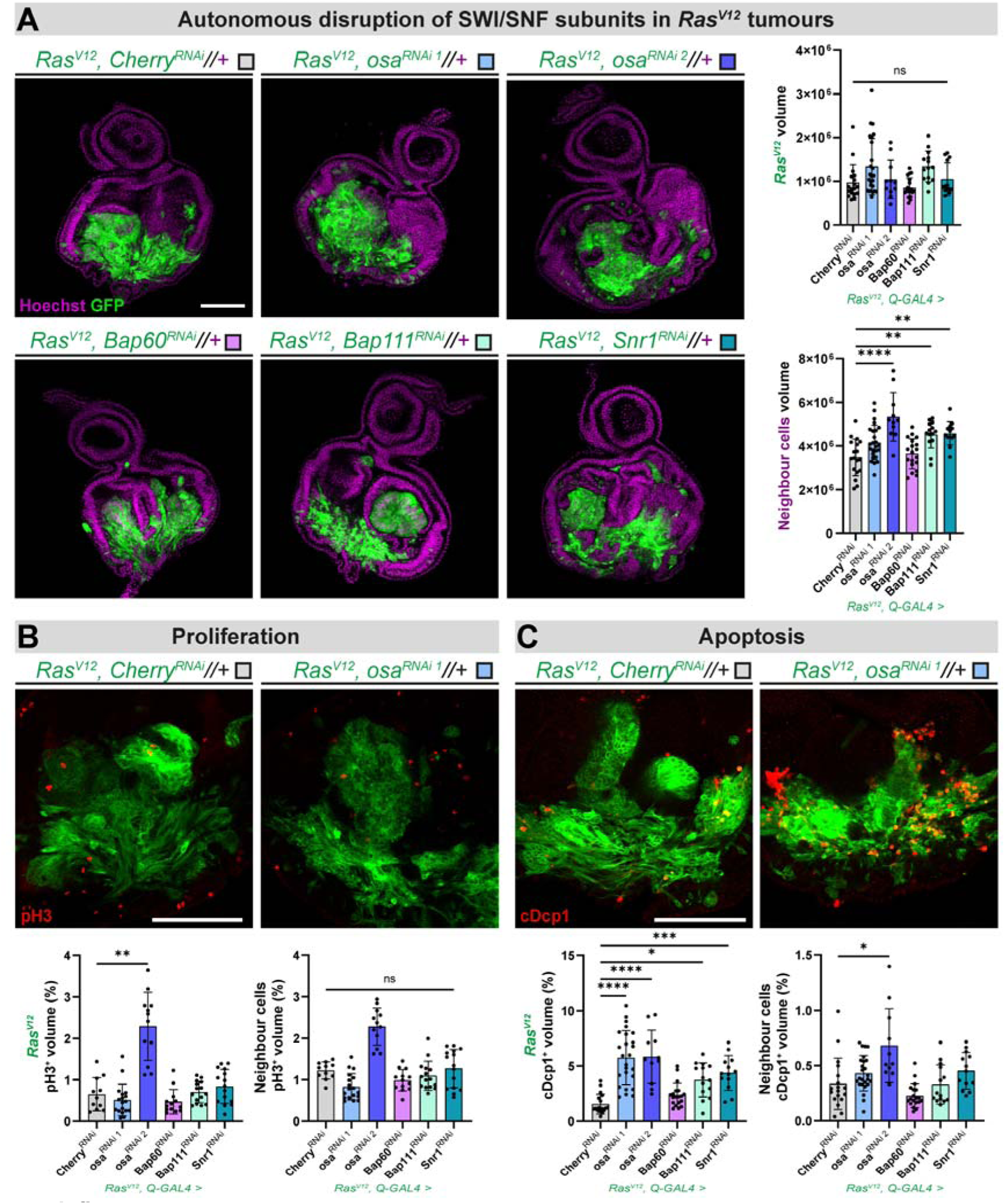
Autonomous disruption of SWI/SNF subunits does not affect *Ras^V12^* clonal growth. (A) Representative confocal images of EADs showing control (*Ras^V12^, Cherry^RNAi^//+*, n=18) and downregulation of different SWI/SNF subunits in *Ras^V12^*cells (*Ras^V12^, osa^RNAi1^//+*, n=25; *Ras^V12^, osa^RNAi2^//+*, n=11; *Ras^V12^, Bap60^RNAi^//+*, n=19; *Ras^V12^, Bap111^RNAi^//+*, n=14; *Ras^V12^, Snr1^RNAi^//+*, n=13). In each condition, *Ras^V12^* cells are labelled with GFP (green), and nuclei are stained with Hoechst (magenta). Quantification of *Ras^V12^* clone volume (top right) and neighbouring tissue volume (bottom right) of distinct EADs is shown (µm^3^). Statistical significance was assessed with Kruskal–Wallis and Dunn’s test for *Ras^V12^* clone volume, and one-way ANOVA was used to test for significance for neighbour tissue volume. Scale bar = 100 µm. (B) Representative confocal images of EADs and quantification of pH3-positive volume fraction within *Ras^V12^*clones and in the neighbouring cells in the genotypes of control (*Ras^V12^, Cherry^RNAi^//+*, n=11) and upon downregulation of different SWI/SNF subunits in *Ras^V12^* cells (*Ras^V12^, osa^RNAi1^//+*, n=18; *Ras^V12^, osa^RNAi2^//+*, n=12; *Ras^V12^, Bap60^RNAi^//+*, n=13; *Ras^V12^, Bap111^RNAi^//+*, n=15; *Ras^V12^, Snr1^RNAi^//+*, n=15). In each condition, *Ras^V12^* cells are labelled with GFP (green), and EAD are stained for pH3 (red). Statistical significance was determined with Kruskal–Wallis and Dunn’s test. Scale bar = 100 µm. (C) Representative confocal images of EADs and quantification of cDcp1-positive volume fraction within within *Ras^V12^* clones and in the neighbouring cells in the genotypes of control (*Ras^V12^, Cherry^RNAi^//+*, n=18) and upon downregulation of different SWI/SNF subunits in *Ras^V12^* cells (*Ras^V12^, osa^RNAi1^//+*, n=25; *Ras^V12^, osa^RNAi2^//+*, n=11; *Ras^V12^, Bap60^RNAi^//+*, n=19; *Ras^V12^, Bap111^RNAi^//+*, n=14; *Ras^V12^, Snr1^RNAi^//+*, n=13). In each condition, *Ras^V12^* cells are labelled with GFP (green), and EAD are stained for cDcp1 (red). Statistical significance was determined with Kruskal–Wallis and Dunn’s test. Scale bar = 100 µm.

## Materials and methods

### Resources table

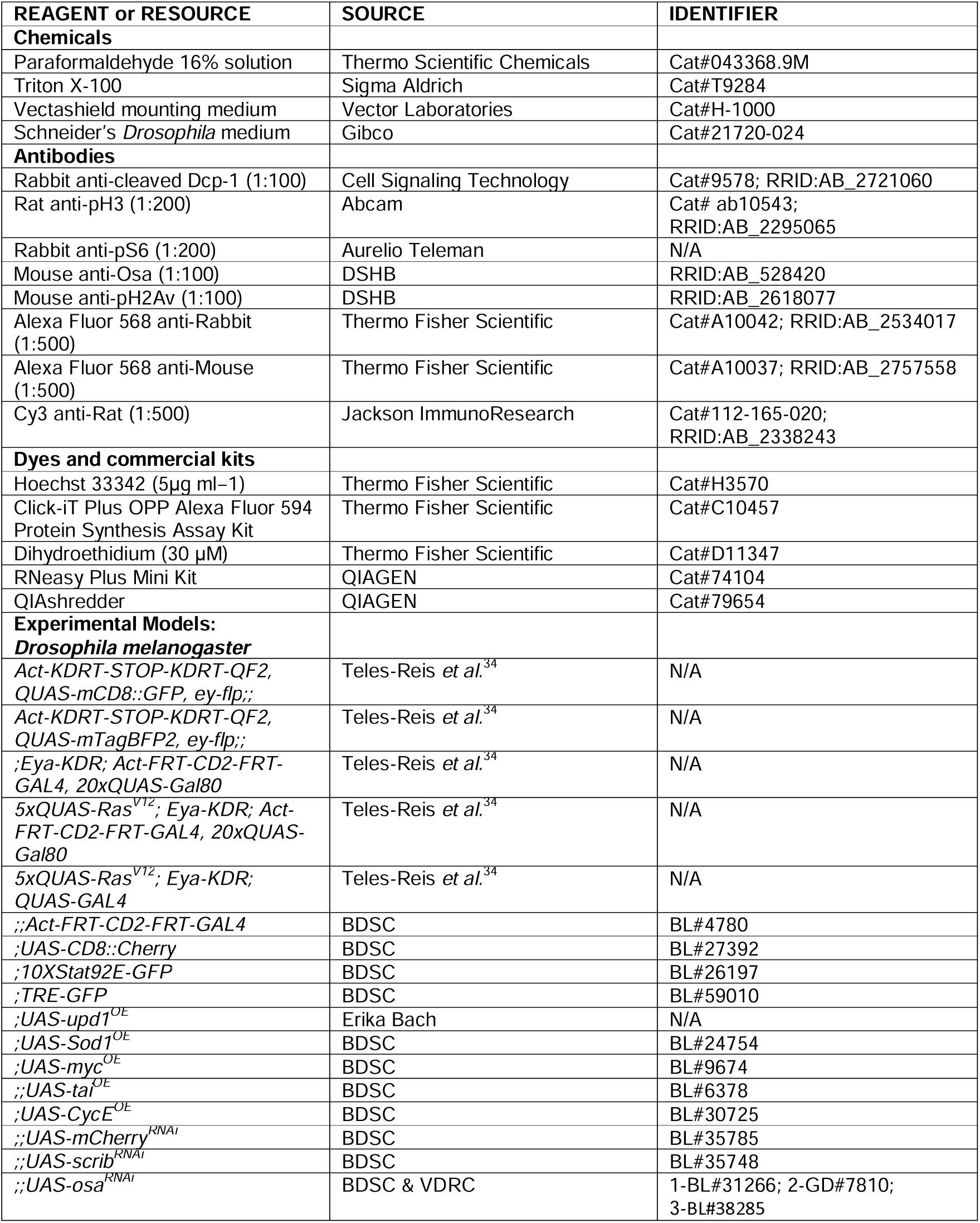
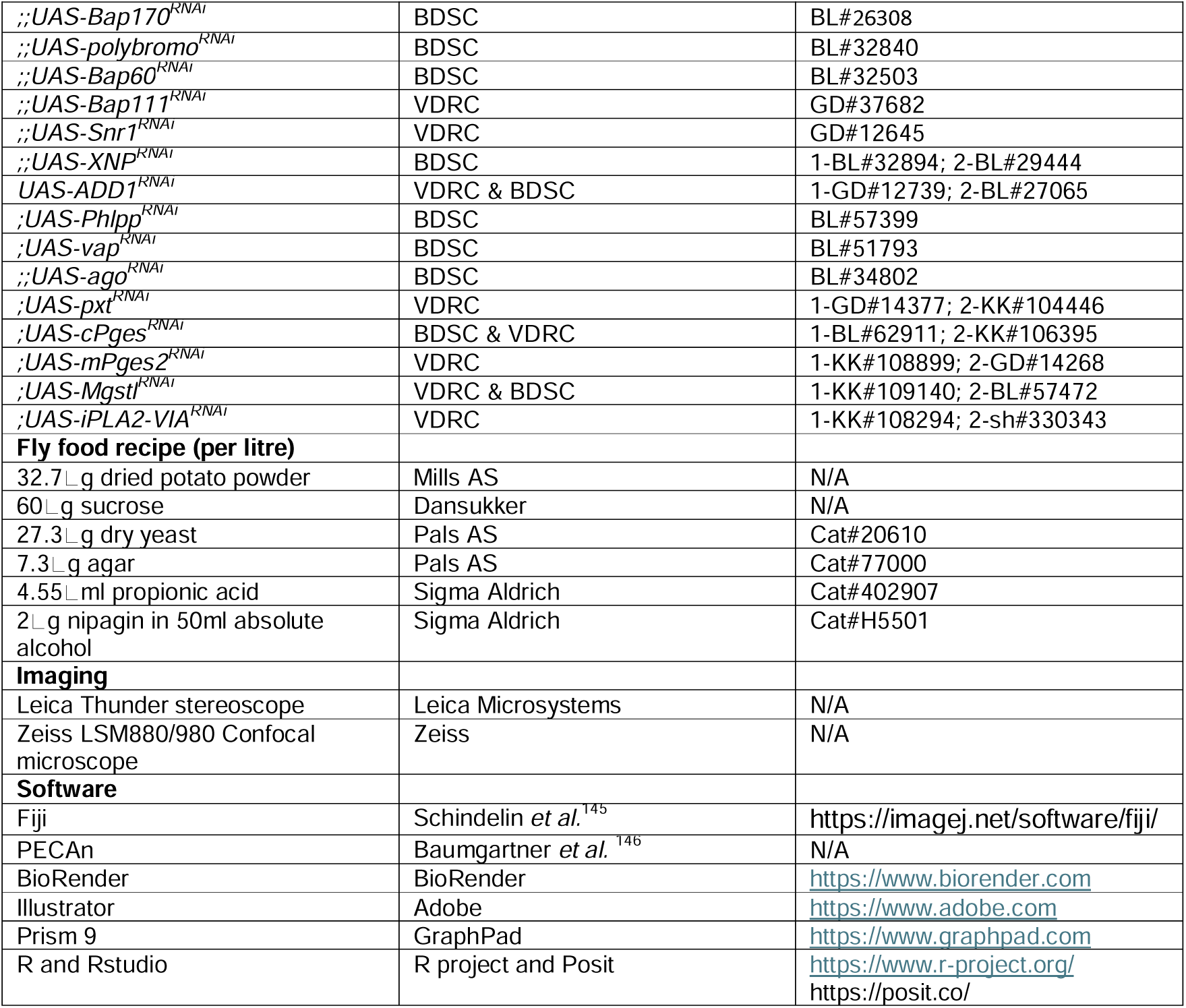

### Method details

#### Computational analysis to identify clinically relevant RAS co-mutated drivers

##### Genes co-mutated with *RAS* mutants

Based on evidence from large collections of human tumour samples, we sought to establish sets of oncogenes and tumour suppressors that were most frequently co-mutated with RAS mutant samples. Specifically, using the TCGAbiolinks (v2.22) package in R^147^, we looked at carcinoma samples across all tissues/sites (i.e. pancancer, n = 7,753) in The Cancer Genome Atlas (TCGA release 31, October 29th 2021), and considered the collection of somatic DNA aberrations detected in these, both short nucleotide variants, insertions/deletions (SNVs/Indels) and gene copy number aberrations. We next devised multiple strategies in order to *i)* define the scope of being a RAS mutant, *ii)* annotate oncogenes/tumour suppressor genes, and *iii)* rank the most highly RAS co-mutated genes established in *ii)*, for which we restricted the set of aberrations to those most likely producing loss-of-function or gain/change-of-function events.

##### *RAS* mutants

*RAS* mutant samples were considered to be those tumour samples with a mutation in *KRAS*/*NRAS*/*HRAS*, and where *i)* the mutation was located in a known mutation hotspot, as found in cancerhotspots.org^148^, and *ii)* the mutation was predicted as a cancer driver mutation^149^.

##### Annotation of oncogenes and tumour suppressor genes

Status of genes as oncogenes and/or tumour suppressors genes were retrieved through a combination of annotations from the CancerMine literature mining resource (v42) and the curated Network of Cancer Genes (NCG v7.0) database^150,151^. Specifically, we required evidence of a gene being a tumour suppressor/oncogene from five or more publications in the biomedical literature (as collected from CancerMine), or at least two publications from CancerMine alongside existing records for the same gene as a tumour suppressor/oncogene in the Network of Cancer Genes (NCG). Status as oncogenes were ignored if a given gene had three times as much support (literature evidence) for a role as a tumour suppressor gene (and vice versa).

##### Loss-of-function versus gain-of-function mutations

We next established TCGA aberrations in oncogenes and tumour suppressors that could be considered as likely loss-of-function (LOF) or gain/change-of-function events (GOF). For SNVs/InDels, we considered LOFs as those mutations with a functional consequence of either stop-gain, frameshift or splice acceptor/donor disruption. For copy number aberrations, we considered homozygous deletions (complete loss of both gene copies, as annotated with a score of −2 by GISTIC2^152^). GOF SNVs/InDels were limited to those that either overlapped with a predicted driver mutation or that were located in a mutation hotspot site^148,149^. For copy number aberrations, GOF events were limited to high-level amplifications (gene level, also by GISTIC2, discrete score of 2).

##### Top *RAS* co-mutated genes in *Drosophila*

By restricting the set of TCGA samples to those annotated as *RAS* mutants, we next summarised, for each oncogene and tumour suppressor candidate, the number of samples mutated, through GOF and LOF, respectively. Genes co-mutated with *RAS* in less than n = 10 samples were ignored. Finally, for the top human *RAS* co-mutated oncogenes and tumour suppressors identified in TCGA using the strategy above, we identified the corresponding fly orthologs through the *babelgene* (v21.4) package in R.

##### Fly husbandry

For all experimental crosses, progenitor flies were allowed to lay eggs for 24 hours in vials with standard potato mash fly food, supplemented with yeast to boost egg-lay. Parents were then removed, and larvae were allowed to develop inside a controlled humidity and temperature (25°C) incubator for 6 days after egg-lay (AEL) before dissection.

##### Drosophila genetics

A complete description of the genotypes per figure can be found in suppl. table 1. Additionally, all the transgenics used in the genetic screen are described in suppl. table 2. The following *Drosophila* stocks were acquired from the Bloomington *Drosophila* stock center (BDSC) *Act-FRT-CD2-FRT-GAL4* (BL#4780), *UAS-CD8::Cherry* (BL#27392), *10Xstat92E-GFP* (BL#26197), *TRE-GFP* (BL#59010), *UAS-Sod1^OE^* (BL#24754), *UAS-myc^OE^* (BL#9674), *UAS-tai^OE^* (BL#6378), *UAS-CycE^OE^* (BL#30725), *UAS-mCherry^RNAi^* (BL#35785), *UAS-scrib^RNAi^* (BL#35748), *UAS-osa^RNAi^* (BL#31266, BL#38285), *UAS-Bap170^RNAi^* (BL#26308), *UAS-polybromo^RNAi^* (BL#32840), *UAS-Bap60^RNAi^* (BL#32503), *UAS-XNP^RNAi^* (BL#32894, BL#29444), *UAS-ADD1^RNAi^* (BL#27065), *UAS-Phlpp^RNAi^* (BL#57399), *UAS-vap^RNAi^* (BL#51793), *UAS-ago^RNAi^* (BL#34802), *UAS-cPges^RNAi^* (BL#62911), *UAS-Mgstl^RNAi^* (BL#57472). Furthermore, the following fly lines were obtained from the Vienna *Drosophila* stock center (VDRC) *UAS-osa^RNAi^* (GD#7810), *UAS-Bap111^RNAi^* (GD#37682), *UAS-Snr1^RNAi^* (GD#12645), *UAS-ADD1^RNAi^* (GD#12739), *UAS-pxt^RNAi^* (GD#14377, KK#104446), *UAS-cPges^RNAi^* (KK#106395), *UAS-mPges2^RNAi^* (KK#108899; GD#14268), *UAS-Mgstl^RNAi^* (KK#109140), *UAS-iPLA2-VIA^RNAi^* (1-KK#108294; 2-sh#330343). *UAS-upd1^OE^* was kindly gifted by Erika Bach.

To independently manipulate two epithelial populations in the eye-antennal disc, we have utilized the EyaHOST system, which was recently described in detail^34^. Briefly, transgenics for misexpression of genes of interest were introduced in the 2^nd^ or 3^rd^ chromosomes of the EyaHOST pickup line *Act-KDRT-STOP-KDRT-QF2, QUAS-mCD8::GFP, ey-flp;;* (GFP^+^ tumour labelling) or *Act-KDRT-STOP-KDRT-QF2, QUAS-mTagBFP2, ey-flp;;* (BFP^+^ tumour labelling), and crossed to different tester lines for clone generation *;Eya-KDR; Act-FRT-CD2-FRT-GAL4, 20xQUAS-Gal80* (WT clones with neighbouring tissue manipulation) or *5xQUAS-Ras^V12^; Eya-KDR; Act-FRT-CD2-FRT-GAL4, 20xQUAS-Gal80* (*Ras^V12^* clones with neighbouring tissue manipulation). Additionally, the following tester line was used *5xQUAS-Ras^V12^; Eya-KDR; QUAS-GAL4* (*Ras^V12^* clones with autonomous genetic manipulation). Because the EyaHOST system relies on transgenes located on the X chromosome, only female larvae could be used in this study. All figures therefore present data exclusively from female larvae and adults (survival experiments).

#### Immunohistochemistry

Day 6 AEL wandering third instar larvae with the correct genotype were collected and briefly washed in 1xPBS. The larvae were then placed in cold 1xPBS on a silicon dissection plate and partially dissected by cutting at midsection and inverting the anterior cuticle to expose the cephalic complex. For each dissected larva, the cephalic complex was transferred into 1.5 mL Eppendorf on ice containing 1xPBS. Dissections were completed under 20 minutes, after which paraformaldehyde was added to a final concentration of 4% to fix the samples for 30 minutes. Samples were then washed three times in PBT (0.5 % Triton X-100 in PBS) and stained with appropriate dyes/antibodies. Concentrations of dyes and antibodies are listed in the key resources table. After staining and final washes with PBT, cephalic complex or individual EADs were mounted with Vectashield in a glass slide with spacers and a coverslip.

The OPP incorporation assay was performed by following the protocol described by Kiparaki and Baker^153^.

The DHE staining was performed as previously described by Pinal *et al.*^102^, with modifications. Larvae were dissected in Schneider’s medium, leaving the EADs attached to the mouth hook. Samples were then incubated in Schneider’s medium containing DHE (30 µM) for 10 minutes, followed by two washes in 1x PBS. Discs were mounted in 1x PBS and imaged immediately using a confocal microscope.

#### Imaging and quantification

Mounted samples were imaged with Zeiss LSM880 or LSM980 confocal microscope and 20x air objective. Whole-tissue z-stacks were acquired for each sample with a z-slice step size of 3 or 4 μm. All microscopy image analysis has been performed with the fiji plugin PECAn^146^, while following the standard workflow. Briefly, compartment specific (clones or surrounding cells) volume quantification was possible through segmentation of tissue autofluorescence and/or nuclear staining, while clones were segmented based on GFP or BFP expression. Fluorescence intensity for STAT-GFP, TRE-GFP, pS6, and OPP, is presented as mean grey value for the compartment indicated. The markers for proliferation (pH3), apoptosis (cDcp1), ROS (DHE), and DNA damage (pH2Av) were measured through compartment specific volume occupancy (%).

#### RNA sequencing

##### Dissection and RNA extraction

Day 6 wandering third instar larvae were washed twice in 1xPBS and dissected in Schneider’s medium. Between 80 and 100 individual eye-antennal discs were dissected within 40 minutes. Every 10 dissected discs were transferred to a 1.5 mL Eppendorf tube on ice containing Schneider’s medium. Prior to transfer, the pipette tip was coated by performing up and down of carcasses to prevent disc sticking and loss.

RNA was extracted using the RNeasy plus mini kit (QIAGEN). Samples were briefly spun down (1-2 seconds) and the Schneider’s medium was removed. Tissue disruption was carried out by adding 200 μl of RLT lysis buffer, followed by mechanical dissociation for 1 minute using a rotor and pestle while the tube was kept on ice. An additional 150 μl of RLT lysis buffer was then added, and the lysate was mixed by pipetting up and down. The sample was further homogenized using a QIAshredder column, and RNA purification was completed according to the RNeasy plus mini kit protocol. Purified RNA was eluted in RNAse-free water, flash frozen in liquid nitrogen, and stored at −80°C. RNA integrity was assessed with Agilent Bioanalyzer system, which showed no evidence of RNA degradation.

##### Library preparation and sequencing

Library preparation and sequencing for the experiments of “*Osa^RNAi^ vs. Cherry^RNAi^*” and “*Bap111^RNAi^ vs. Cherry^RNAi^*” was performed by the Genomics core facility at the Oslo University Hospital (OUH) with Illumina stranded mRNA preparation and NextSeq500 paired-end 2×75 bp. The “*Ras^V12^//Osa^RNAi^ vs. Ras^V12^// Cherry^RNAi^*” was processed by Novogene with mRNA poly A enrichment library preparation and NovaSeq X Plus paired-end 2×150bp sequencing.

#### Transcriptome computational analysis

All RNA sequencing computational analyses were conducted using the nf-core RNA-seq pipeline version 3.14.0 (https://nf-co.re/rnaseq/3.14.0/). This pipeline ensures reproducibility in processing RNAseq data by providing a standardized workflow. Various tools were employed within the pipeline, each serving distinct functions throughout the RNAseq data processing stages.

##### Preprocessing and quality control

Adaptor trimming and filtering of low-quality reads were performed using Fastp. FastQC was used to check for base quality scores, GC content, and the presence of adapter sequences. RSeQC was further used to evaluate read distribution and coverage.

##### Alignment, deduplication, and quantification

Sequencing reads were aligned to the reference genome of *Drosophila melanogaster* (BDGP6.46) using the STAR aligner. The aligned reads were then sorted and indexed using SAMtools. To mitigate the effects of PCR duplicates, UMI-tools was used for UMI-based deduplication, and Picard MarkDuplicates was applied to mark duplicate reads in the dataset. Finally, Salmon was used to perform transcript quantification, calculating Transcripts Per Million (TPM) counts.

##### Differential Expression Analysis Tools

For differential expression analysis, DESeq2 package was used in R. DESeq2 compares gene expression profiles across conditions and provides estimates of dispersion and log2 fold change (Log2FC). Differentially expressed genes (DEGs) were defined with a |Log_JFC| threshold ≥ 0.58 and an adjusted p-value threshold of ≤ 0.05.

##### Visualization

Visualization of the results of differential gene expression analysis was carried out using R. The ggplot2 package was used for creating volcano plots, and the ggVennDiagram package allowed the generation of venn diagrams to study the intersection of differentially expressed genes between datasets.

#### Statistical analysis

Statistical analysis was performed using GraphPad prism 9. Datasets were tested for normality using the Shapiro-Wilk test. The suitable parametric or non-parametric test were applied to test for statistical significance between groups. The test used can be found in the figure legends. All statistical tests performed were two-sided. Significance is indicated as: ns, *P* ≥ 0.05; *, *P* < 0.05; **, *P* < 0.01; ***, *P* < 0.001; ****, *P* < 0.0001. Error bars represent standard deviation. At least two replicates have been performed in all experiments. Detailed results for the statistical analysis performed for each figure is provided in the Source Data file.

## Resource and data availability

Source data for each figure is provided in the Source Data file. For requests, contact Tor Erik Rusten.

## Acknowledgements

We would like to thank Aurelio Teleman, Erika Bach, the Developmental Studies Hybridoma Bank, the Bloomington *Drosophila* Stock Centre, the Transgenic RNAi Project at Harvard Medical School, the Vienna *Drosophila* Stock Centre, for *Drosophila* lines and antibodies. We additionally thank the support from the core facility for Advanced Light Microscopy and the Genomics core facility at Oslo University Hospital.

This project was funded by the Norwegian Research Council Toppforsk and Center of Excellence, grants #262652 and #276070 & The Norwegian Cancer Society #247130. A.M.D was funded by InFLAMES Flagship Programme of the Academy of Finland, grant #337531.

## Author contributions

Conceptualization: J.T.R., T.E.R.; Methodology: J.T.R., C.D., A.J., M.E.B.; Investigation: J.T.R., C.D., M.G.A., P.R.D., M.D., D.L., C.G., S.N., A.S., A.M.D., V.R.; Data curation: J.T.R., S.N., A.S.; Formal analysis: J.T.R., S.N., A.S.; Software: J.T.R., S.N., A.S., M.E.B.; Visualization: J.T.R.; Writing – original draft: J.T.R., T.E.R.; Writing – review & editing: J.T.R., T.E.R.; Supervision: T.E.R.; Funding acquisition: T.E.R.

