## Supplemental table 1 for "Interclonal cooperation and suppression shape early Ras-driven tumour growth"

<table>
<tr><th rowspan="2">Figure</th><th colspan="3">Progenitor genotype</th></tr>
<tr><th>♀</th><th>x</th><th>♂</th></tr>
<tr><td rowspan="5">1B</td><td>Actin>|STOP>|QF2, QUAS-CD8::GFP, ey-FLP;;</td><td>x</td><td>5xQUAS-RasV12; eya-KD; Act>STOP>GAL4, 20xQUAS-GAL80</td></tr>
<tr><td>Actin>|STOP>|QF2, QUAS-CD8::GFP, ey-FLP;;UAS-CherryRNAi</td><td>x</td><td>5xQUAS-RasV12; eya-KD; Act>STOP>GAL4, 20xQUAS-GAL80</td></tr>
<tr><td>Actin>|STOP>|QF2, QUAS-CD8::GFP, ey-FLP;UAS-CherryOE</td><td>x</td><td>5xQUAS-RasV12; eya-KD; Act>STOP>GAL4, 20xQUAS-GAL80</td></tr>
<tr><td>Actin>|STOP>|QF2, QUAS-CD8::GFP, ey-FLP;;UAS-scribRNAi</td><td>x</td><td>5xQUAS-RasV12; eya-KD; Act>STOP>GAL4, 20xQUAS-GAL80</td></tr>
<tr><td>Actin>|STOP>|QF2, QUAS-CD8::GFP, ey-FLP;UAS-upd1OE</td><td>x</td><td>5xQUAS-RasV12; eya-KD; Act>STOP>GAL4, 20xQUAS-GAL80</td></tr>
<tr><td rowspan="6">2A</td><td>Actin>|STOP>|QF2, QUAS-CD8::GFP, ey-FLP;;UAS-CherryRNAi</td><td>x</td><td>5xQUAS-RasV12; eya-KD; Act>STOP>GAL4, 20xQUAS-GAL80</td></tr>
<tr><td>Actin>|STOP>|QF2, QUAS-CD8::GFP, ey-FLP;;UAS-osaRNAi 3</td><td>x</td><td>5xQUAS-RasV12; eya-KD; Act>STOP>GAL4, 20xQUAS-GAL80</td></tr>
<tr><td>Actin>|STOP>|QF2, QUAS-CD8::GFP, ey-FLP;;UAS-Bap170RNAi</td><td>x</td><td>5xQUAS-RasV12; eya-KD; Act>STOP>GAL4, 20xQUAS-GAL80</td></tr>
<tr><td>Actin>|STOP>|QF2, QUAS-CD8::GFP, ey-FLP;;UAS-polybromoRNAi</td><td>x</td><td>5xQUAS-RasV12; eya-KD; Act>STOP>GAL4, 20xQUAS-GAL80</td></tr>
<tr><td>Actin>|STOP>|QF2, QUAS-CD8::GFP, ey-FLP;;UAS-XNPRNAi 1</td><td>x</td><td>5xQUAS-RasV12; eya-KD; Act>STOP>GAL4, 20xQUAS-GAL80</td></tr>
<tr><td>Actin>|STOP>|QF2, QUAS-CD8::GFP, ey-FLP;UAS-PhlppRNAi</td><td>x</td><td>5xQUAS-RasV12; eya-KD; Act>STOP>GAL4, 20xQUAS-GAL80</td></tr>
<tr><td rowspan="6">3A</td><td>Actin>|STOP>|QF2, QUAS-CD8::GFP, ey-FLP;;UAS-CherryRNAi</td><td>x</td><td>5xQUAS-RasV12; eya-KD; Act>STOP>GAL4, 20xQUAS-GAL80</td></tr>
<tr><td>Actin>|STOP>|QF2, QUAS-CD8::GFP, ey-FLP;UAS-vapRNAi</td><td>x</td><td>5xQUAS-RasV12; eya-KD; Act>STOP>GAL4, 20xQUAS-GAL80</td></tr>
<tr><td>Actin>|STOP>|QF2, QUAS-CD8::GFP, ey-FLP;;UAS-agoRNAi</td><td>x</td><td>5xQUAS-RasV12; eya-KD; Act>STOP>GAL4, 20xQUAS-GAL80</td></tr>
<tr><td>Actin>|STOP>|QF2, QUAS-CD8::GFP, ey-FLP;UAS-CherryOE</td><td>x</td><td>5xQUAS-RasV12; eya-KD; Act>STOP>GAL4, 20xQUAS-GAL80</td></tr>
<tr><td>Actin>|STOP>|QF2, QUAS-CD8::GFP, ey-FLP;;UAS-taiOE</td><td>x</td><td>5xQUAS-RasV12; eya-KD; Act>STOP>GAL4, 20xQUAS-GAL80</td></tr>
<tr><td>Actin>|STOP>|QF2, QUAS-CD8::GFP, ey-FLP;UAS-mycOE</td><td>x</td><td>5xQUAS-RasV12; eya-KD; Act>STOP>GAL4, 20xQUAS-GAL80</td></tr>
<tr><td rowspan="2">3B</td><td>Actin>|STOP>|QF2, QUAS-CD8::GFP, ey-FLP;;</td><td>x</td><td>5xQUAS-RasV12; eya-KD; Act>STOP>GAL4, 20xQUAS-GAL80</td></tr>
<tr><td>Actin>|STOP>|QF2, QUAS-CD8::GFP, ey-FLP;UAS-CycEOE</td><td>x</td><td>5xQUAS-RasV12; eya-KD; Act>STOP>GAL4, 20xQUAS-GAL80</td></tr>
<tr><td rowspan="8">3C</td><td>Actin>|STOP>|QF2, QUAS-CD8::GFP, ey-FLP;;UAS-CherryRNAi</td><td>x</td><td>5xQUAS-RasV12; eya-KD; Act>STOP>GAL4, 20xQUAS-GAL80</td></tr>
<tr><td>Actin>|STOP>|QF2, QUAS-CD8::GFP, ey-FLP;UAS-vapRNAi</td><td>x</td><td>5xQUAS-RasV12; eya-KD; Act>STOP>GAL4, 20xQUAS-GAL80</td></tr>
<tr><td>Actin>|STOP>|QF2, QUAS-CD8::GFP, ey-FLP;;UAS-agoRNAi</td><td>x</td><td>5xQUAS-RasV12; eya-KD; Act>STOP>GAL4, 20xQUAS-GAL80</td></tr>
<tr><td>Actin>|STOP>|QF2, QUAS-CD8::GFP, ey-FLP;;</td><td>x</td><td>5xQUAS-RasV12; eya-KD; Act>STOP>GAL4, 20xQUAS-GAL80</td></tr>
<tr><td>Actin>|STOP>|QF2, QUAS-CD8::GFP, ey-FLP;UAS-CycEOE</td><td>x</td><td>5xQUAS-RasV12; eya-KD; Act>STOP>GAL4, 20xQUAS-GAL80</td></tr>
<tr><td>Actin>|STOP>|QF2, QUAS-CD8::GFP, ey-FLP;UAS-CherryOE</td><td>x</td><td>5xQUAS-RasV12; eya-KD; Act>STOP>GAL4, 20xQUAS-GAL80</td></tr>
<tr><td>Actin>|STOP>|QF2, QUAS-CD8::GFP, ey-FLP;;UAS-taiOE</td><td>x</td><td>5xQUAS-RasV12; eya-KD; Act>STOP>GAL4, 20xQUAS-GAL80</td></tr>
<tr><td>Actin>|STOP>|QF2, QUAS-CD8::GFP, ey-FLP;UAS-mycOE</td><td>x</td><td>5xQUAS-RasV12; eya-KD; Act>STOP>GAL4, 20xQUAS-GAL80</td></tr>
<tr><td rowspan="6">4B</td><td>Actin>|STOP>|QF2, QUAS-CD8::GFP, ey-FLP;;UAS-CherryRNAi</td><td>x</td><td>5xQUAS-RasV12; eya-KD; Act>STOP>GAL4, 20xQUAS-GAL80</td></tr>
<tr><td>Actin>|STOP>|QF2, QUAS-CD8::GFP, ey-FLP;;UAS-osaRNAi 1</td><td>x</td><td>5xQUAS-RasV12; eya-KD; Act>STOP>GAL4, 20xQUAS-GAL80</td></tr>
<tr><td>Actin>|STOP>|QF2, QUAS-CD8::GFP, ey-FLP;;UAS-osaRNAi 2</td><td>x</td><td>5xQUAS-RasV12; eya-KD; Act>STOP>GAL4, 20xQUAS-GAL80</td></tr>
<tr><td>Actin>|STOP>|QF2, QUAS-CD8::GFP, ey-FLP;;UAS-Bap60RNAi</td><td>x</td><td>5xQUAS-RasV12; eya-KD; Act>STOP>GAL4, 20xQUAS-GAL80</td></tr>
<tr><td>Actin>|STOP>|QF2, QUAS-CD8::GFP, ey-FLP;;UAS-Bap111RNAi</td><td>x</td><td>5xQUAS-RasV12; eya-KD; Act>STOP>GAL4, 20xQUAS-GAL80</td></tr>
<tr><td>Actin>|STOP>|QF2, QUAS-CD8::GFP, ey-FLP;;UAS-Snr1RNAi</td><td>x</td><td>5xQUAS-RasV12; eya-KD; Act>STOP>GAL4, 20xQUAS-GAL80</td></tr>
<tr><td rowspan="2">4C</td><td>Actin>|STOP>|QF2, QUAS-CD8::GFP, ey-FLP;;UAS-CherryRNAi</td><td>x</td><td>5xQUAS-RasV12; eya-KD; Act>STOP>GAL4, 20xQUAS-GAL80</td></tr>
<tr><td>Actin>|STOP>|QF2, QUAS-CD8::GFP, ey-FLP;;UAS-osaRNAi 1</td><td>x</td><td>5xQUAS-RasV12; eya-KD; Act>STOP>GAL4, 20xQUAS-GAL80</td></tr>
<tr><td rowspan="2">4D</td><td>Actin>|STOP>|QF2, QUAS-CD8::GFP, ey-FLP;;UAS-CherryRNAi</td><td>x</td><td>5xQUAS-RasV12; eya-KD; Act>STOP>GAL4, 20xQUAS-GAL80</td></tr>
<tr><td>Actin>|STOP>|QF2, QUAS-CD8::GFP, ey-FLP;;UAS-osaRNAi 1</td><td>x</td><td>5xQUAS-RasV12; eya-KD; Act>STOP>GAL4, 20xQUAS-GAL80</td></tr>
<tr><td rowspan="6">5A,B,C</td><td>Actin>|STOP>|QF2, QUAS-CD8::GFP, ey-FLP;;UAS-CherryRNAi</td><td>x</td><td>;;Act>STOP>GAL4</td></tr>
<tr><td>Actin>|STOP>|QF2, QUAS-CD8::GFP, ey-FLP;;UAS-Bap111RNAi</td><td>x</td><td>;;Act>STOP>GAL4</td></tr>
<tr><td>Actin>|STOP>|QF2, QUAS-CD8::GFP, ey-FLP;;UAS-CherryRNAi</td><td>x</td><td>;;Act>STOP>GAL4</td></tr>
<tr><td>Actin>|STOP>|QF2, QUAS-CD8::GFP, ey-FLP;;UAS-osaRNAi 1</td><td>x</td><td>;;Act>STOP>GAL4</td></tr>
<tr><td>Actin>|STOP>|QF2, QUAS-CD8::GFP, ey-FLP;;UAS-CherryRNAi</td><td>x</td><td>5xQUAS-RasV12; eya-KD; Act>STOP>GAL4, 20xQUAS-GAL80</td></tr>
<tr><td>Actin>|STOP>|QF2, QUAS-CD8::GFP, ey-FLP;;UAS-osaRNAi 1</td><td>x</td><td>5xQUAS-RasV12; eya-KD; Act>STOP>GAL4, 20xQUAS-GAL80</td></tr>
<tr><td rowspan="6">6A</td><td>Actin>|STOP>|QF2, QUAS-mTagBFP2, ey-FLP;STAT-GFP;+</td><td>x</td><td>; eya-KD; Act>STOP>GAL4, 20xQUAS-GAL80</td></tr>
<tr><td>Actin>|STOP>|QF2, QUAS-mTagBFP2, ey-FLP;STAT-GFP;+</td><td>x</td><td>5xQUAS-RasV12; eya-KD; Act>STOP>GAL4, 20xQUAS-GAL80</td></tr>
<tr><td>Actin>|STOP>|QF2, QUAS-mTagBFP2, ey-FLP;STAT-GFP;UAS-osaRNAi 1</td><td>x</td><td>5xQUAS-RasV12; eya-KD; Act>STOP>GAL4, 20xQUAS-GAL80</td></tr>
<tr><td>Actin>|STOP>|QF2, QUAS-mTagBFP2, ey-FLP;TRE-GFP;+</td><td>x</td><td>; eya-KD; Act>STOP>GAL4, 20xQUAS-GAL80</td></tr>
<tr><td>Actin>|STOP>|QF2, QUAS-mTagBFP2, ey-FLP;TRE-GFP;+</td><td>x</td><td>5xQUAS-RasV12; eya-KD; Act>STOP>GAL4, 20xQUAS-GAL80</td></tr>
<tr><td>Actin>|STOP>|QF2, QUAS-mTagBFP2, ey-FLP;TRE-GFP;UAS-osaRNAi 1</td><td>x</td><td>5xQUAS-RasV12; eya-KD; Act>STOP>GAL4, 20xQUAS-GAL80</td></tr>
<tr><td rowspan="2">6B,C</td><td>Actin>|STOP>|QF2, QUAS-CD8::GFP, ey-FLP;;UAS-CherryRNAi</td><td>x</td><td>5xQUAS-RasV12; eya-KD; Act>STOP>GAL4, 20xQUAS-GAL80</td></tr>
<tr><td>Actin>|STOP>|QF2, QUAS-CD8::GFP, ey-FLP;;UAS-osaRNAi 1</td><td>x</td><td>5xQUAS-RasV12; eya-KD; Act>STOP>GAL4, 20xQUAS-GAL80</td></tr>
<tr><td rowspan="9">6D</td><td>Actin>|STOP>|QF2, QUAS-CD8::GFP, ey-FLP;;UAS-CherryRNAi</td><td>x</td><td>5xQUAS-RasV12; eya-KD; Act>STOP>GAL4, 20xQUAS-GAL80</td></tr>
<tr><td>Actin>|STOP>|QF2, QUAS-CD8::GFP, ey-FLP;;UAS-osaRNAi 1</td><td>x</td><td>5xQUAS-RasV12; eya-KD; Act>STOP>GAL4, 20xQUAS-GAL80</td></tr>
<tr><td>Actin>|STOP>|QF2, QUAS-CD8::GFP, ey-FLP;;UAS-Bap60RNAi</td><td>x</td><td>5xQUAS-RasV12; eya-KD; Act>STOP>GAL4, 20xQUAS-GAL80</td></tr>
<tr><td>Actin>|STOP>|QF2, QUAS-CD8::GFP, ey-FLP;;UAS-Snr1RNAi</td><td>x</td><td>5xQUAS-RasV12; eya-KD; Act>STOP>GAL4, 20xQUAS-GAL80</td></tr>
<tr><td>Actin>|STOP>|QF2, QUAS-CD8::GFP, ey-FLP;;UAS-CherryRNAi</td><td>x</td><td>5xQUAS-RasV12; eya-KD; Act>STOP>GAL4, 20xQUAS-GAL80</td></tr>
<tr><td>Actin>|STOP>|QF2, QUAS-CD8::GFP, ey-FLP;;UAS-osaRNAi 1</td><td>x</td><td>5xQUAS-RasV12; eya-KD; Act>STOP>GAL4, 20xQUAS-GAL80</td></tr>
<tr><td>Actin>|STOP>|QF2, QUAS-CD8::GFP, ey-FLP;;UAS-Bap60RNAi</td><td>x</td><td>5xQUAS-RasV12; eya-KD; Act>STOP>GAL4, 20xQUAS-GAL80</td></tr>
<tr><td>Actin>|STOP>|QF2, QUAS-CD8::GFP, ey-FLP;;UAS-Bap111RNAi</td><td>x</td><td>5xQUAS-RasV12; eya-KD; Act>STOP>GAL4, 20xQUAS-GAL80</td></tr>
<tr><td>Actin>|STOP>|QF2, QUAS-CD8::GFP, ey-FLP;;UAS-Snr1RNAi</td><td>x</td><td>5xQUAS-RasV12; eya-KD; Act>STOP>GAL4, 20xQUAS-GAL80</td></tr>
<tr><td rowspan="2">7B</td><td>Actin>|STOP>|QF2, QUAS-CD8::GFP, ey-FLP;;UAS-osaRNAi 1</td><td>x</td><td>5xQUAS-RasV12; eya-KD; Act>STOP>GAL4, 20xQUAS-GAL80</td></tr>
<tr><td>Actin>|STOP>|QF2, QUAS-CD8::GFP, ey-FLP;UAS-Sod1OE;UAS-osaRNAi 1</td><td>x</td><td>5xQUAS-RasV12; eya-KD; Act>STOP>GAL4, 20xQUAS-GAL80</td></tr>
<tr><td rowspan="11">7D</td><td>Actin>|STOP>|QF2, QUAS-CD8::GFP, ey-FLP;;UAS-osaRNAi 1</td><td>x</td><td>5xQUAS-RasV12; eya-KD; Act>STOP>GAL4, 20xQUAS-GAL80</td></tr>
<tr><td>Actin>|STOP>|QF2, QUAS-CD8::GFP, ey-FLP;UAS-pxtRNAi 1;UAS-osaRNAi 1</td><td>x</td><td>5xQUAS-RasV12; eya-KD; Act>STOP>GAL4, 20xQUAS-GAL80</td></tr>
<tr><td>Actin>|STOP>|QF2, QUAS-CD8::GFP, ey-FLP;UAS-pxtRNAi 2;UAS-osaRNAi 1</td><td>x</td><td>5xQUAS-RasV12; eya-KD; Act>STOP>GAL4, 20xQUAS-GAL80</td></tr>
<tr><td>Actin>|STOP>|QF2, QUAS-CD8::GFP, ey-FLP;UAS-cPgesRNAi 1;UAS-osaRNAi 1</td><td>x</td><td>5xQUAS-RasV12; eya-KD; Act>STOP>GAL4, 20xQUAS-GAL80</td></tr>
<tr><td>Actin>|STOP>|QF2, QUAS-CD8::GFP, ey-FLP;UAS-cPgesRNAi 2;UAS-osaRNAi 1</td><td>x</td><td>5xQUAS-RasV12; eya-KD; Act>STOP>GAL4, 20xQUAS-GAL80</td></tr>
<tr><td>Actin>|STOP>|QF2, QUAS-CD8::GFP, ey-FLP;UAS-mPges2RNAi 1;UAS-osaRNAi 1</td><td>x</td><td>5xQUAS-RasV12; eya-KD; Act>STOP>GAL4, 20xQUAS-GAL80</td></tr>
<tr><td>Actin>|STOP>|QF2, QUAS-CD8::GFP, ey-FLP;UAS-mPges2RNAi 2;UAS-osaRNAi 1</td><td>x</td><td>5xQUAS-RasV12; eya-KD; Act>STOP>GAL4, 20xQUAS-GAL80</td></tr>
<tr><td>Actin>|STOP>|QF2, QUAS-CD8::GFP, ey-FLP;UAS-MgstlRNAi 1;UAS-osaRNAi 1</td><td>x</td><td>5xQUAS-RasV12; eya-KD; Act>STOP>GAL4, 20xQUAS-GAL80</td></tr>
<tr><td>Actin>|STOP>|QF2, QUAS-CD8::GFP, ey-FLP;UAS-MgstlRNAi 2;UAS-osaRNAi 1</td><td>x</td><td>5xQUAS-RasV12; eya-KD; Act>STOP>GAL4, 20xQUAS-GAL80</td></tr>
<tr><td>Actin>|STOP>|QF2, QUAS-CD8::GFP, ey-FLP;UAS-iPLA2-VIARNAi 1;UAS-osaRNAi 1</td><td>x</td><td>5xQUAS-RasV12; eya-KD; Act>STOP>GAL4, 20xQUAS-GAL80</td></tr>
<tr><td>Actin>|STOP>|QF2, QUAS-CD8::GFP, ey-FLP;UAS-iPLA2-VIARNAi 2;UAS-osaRNAi 1</td><td>x</td><td>5xQUAS-RasV12; eya-KD; Act>STOP>GAL4, 20xQUAS-GAL80</td></tr>
<tr><td rowspan="4">S1A</td><td>Actin>|STOP>|QF2, QUAS-CD8::GFP, ey-FLP;;</td><td>x</td><td>5xQUAS-RasV12; eya-KD; Act>STOP>GAL4, 20xQUAS-GAL80</td></tr>
<tr><td>Actin>|STOP>|QF2, QUAS-CD8::GFP, ey-FLP;;UAS-XNPRNAi 2</td><td>x</td><td>5xQUAS-RasV12; eya-KD; Act>STOP>GAL4, 20xQUAS-GAL80</td></tr>
<tr><td>Actin>|STOP>|QF2, QUAS-CD8::GFP, ey-FLP;UAS-ADD1RNAi 1</td><td>x</td><td>5xQUAS-RasV12; eya-KD; Act>STOP>GAL4, 20xQUAS-GAL80</td></tr>
<tr><td>Actin>|STOP>|QF2, QUAS-CD8::GFP, ey-FLP;;UAS-ADD1RNAi 2</td><td>x</td><td>5xQUAS-RasV12; eya-KD; Act>STOP>GAL4, 20xQUAS-GAL80</td></tr>
<tr><td rowspan="3">S2A</td><td>Actin>|STOP>|QF2, QUAS-CD8::GFP, ey-FLP;;UAS-CherryRNAi</td><td>x</td><td>5xQUAS-RasV12; eya-KD; Act>STOP>GAL4, 20xQUAS-GAL80</td></tr>
<tr><td>Actin>|STOP>|QF2, QUAS-CD8::GFP, ey-FLP;;UAS-osaRNAi 1</td><td>x</td><td>5xQUAS-RasV12; eya-KD; Act>STOP>GAL4, 20xQUAS-GAL80</td></tr>
<tr><td>Actin>|STOP>|QF2, QUAS-CD8::GFP, ey-FLP;;UAS-osaRNAi 2</td><td>x</td><td>5xQUAS-RasV12; eya-KD; Act>STOP>GAL4, 20xQUAS-GAL80</td></tr>
<tr><td rowspan="6">S3A</td><td>Actin>|STOP>|QF2, QUAS-CD8::GFP, ey-FLP;;UAS-CherryRNAi</td><td>x</td><td>5xQUAS-RasV12; eya-KD; QUAS-GAL4</td></tr>
<tr><td>Actin>|STOP>|QF2, QUAS-CD8::GFP, ey-FLP;;UAS-osaRNAi 1</td><td>x</td><td>5xQUAS-RasV12; eya-KD; QUAS-GAL4</td></tr>
<tr><td>Actin>|STOP>|QF2, QUAS-CD8::GFP, ey-FLP;;UAS-osaRNAi 2</td><td>x</td><td>5xQUAS-RasV12; eya-KD; QUAS-GAL4</td></tr>
<tr><td>Actin>|STOP>|QF2, QUAS-CD8::GFP, ey-FLP;;UAS-Bap60RNAi</td><td>x</td><td>5xQUAS-RasV12; eya-KD; QUAS-GAL4</td></tr>
<tr><td>Actin>|STOP>|QF2, QUAS-CD8::GFP, ey-FLP;;UAS-Bap111RNAi</td><td>x</td><td>5xQUAS-RasV12; eya-KD; QUAS-GAL4</td></tr>
<tr><td>Actin>|STOP>|QF2, QUAS-CD8::GFP, ey-FLP;;UAS-Snr1RNAi</td><td>x</td><td>5xQUAS-RasV12; eya-KD; QUAS-GAL4</td></tr>
<tr><td rowspan="2">S3B</td><td>Actin>|STOP>|QF2, QUAS-CD8::GFP, ey-FLP;;UAS-CherryRNAi</td><td>x</td><td>5xQUAS-RasV12; eya-KD; QUAS-GAL4</td></tr>
<tr><td>Actin>|STOP>|QF2, QUAS-CD8::GFP, ey-FLP;;UAS-osaRNAi 1</td><td>x</td><td>5xQUAS-RasV12; eya-KD; QUAS-GAL4</td></tr>
<tr><td rowspan="2">S3C</td><td>Actin>|STOP>|QF2, QUAS-CD8::GFP, ey-FLP;;UAS-CherryRNAi</td><td>x</td><td>5xQUAS-RasV12; eya-KD; QUAS-GAL4</td></tr>
<tr><td>Actin>|STOP>|QF2, QUAS-CD8::GFP, ey-FLP;;UAS-osaRNAi 1</td><td>x</td><td>5xQUAS-RasV12; eya-KD; QUAS-GAL4</td></tr>
</table>
