## Supplemental table 2 for "Interclonal cooperation and suppression shape early Ras-driven tumour growth"

### LOF-Loss Of Function

| Fly_gene | FlyBaseID | Human_ge | Frequency | Mut_Type_ | Mut_Type_ | Role | Transgenic | Type transgenic |
|---|---|---|---|---|---|---|---|---|
| Apc; Apc2 | FBgn00155 | APC | 225 | snv_indel | Hom_del | TSg | 34869 | Val20 attP2 |
| p53 | FBgn0039C | TP53 | 127 | snv_indel | Hom_del | TSg | 41720 | Val20 attP2 |
| osa | FBgn02618 | ARID1A | 116 | snv_indel | Hom_del | TSg | 38285 | Val20 attP40 |
| Pten | FBgn00263 | PTEN | 113 | snv_indel | Hom_del | TSg | 33643 | Val20 attP2 |
| Mtap | FBgn00342 | MTAP | 84 | no | Hom_del | TSg | 56010 | Val20 attP40 |
| Med | FBgn0011€ | SMAD4 | 74 | snv_indel | Hom_del | TSg | 31928 | Val10 attP2 |
| LRP1; mgl | FBgn0053C | LRP1B | 56 | snv_indel | Hom_del | TSg | 44579 | Val20 attP2 |
| Wwox | FBgn00319 | WWOX | 56 | snv_indel | Hom_del | TSg | 51747 | Val20 attP2 |
| Lpt; trr | FBgn0263€ | KMT2D | 52 | snv_indel | no | Onc/TSg | 25994 | Val10 attP2 |
| kug | FBgn02615 | FAT1 | 42 | snv_indel | Hom_del | TSg | 40888 | Val20 attP40 |
| Lar | FBgn00004 | PTPRD | 41 | snv_indel | Hom_del | TSg | 34965 | Val20 attP2 |
| tefu | FBgn0045C | ATM | 39 | snv_indel | Hom_del | TSg | 44073 | Val20 attP40 |
| Smox | FBgn00258 | SMAD2 | 37 | snv_indel | Hom_del | TSg | 41670 | Val20 attP40 |
| ago | FBgn00411 | FBXW7 | 36 | snv_indel | Hom_del | TSg | 34802 | Val20 attP2 |
| Ostgamma | FBgn0032C | TUSC3 | 34 | no | Hom_del | Onc/TSg | 34638 | Val20 attP2 |
| trr | FBgn00235 | KMT2C | 33 | snv_indel | no | TSg | 36916 | Val20 attP2 |
| CG15365 | FBgn0030C | LZTS1 | 33 | snv_indel | Hom_del | TSg | 36856 | Val20 attP2 |
| MCPH1 | FBgn02609 | MCPH1 | 33 | snv_indel | Hom_del | TSg | 38244 | Val20 attP40 |
| zfh2 | FBgn0004€ | ZFHX3 | 32 | snv_indel | Hom_del | TSg | 50643 | Val20 attP2 |
| RhoBTB | FBgn00369 | RHOBTB2 | 31 | no | Hom_del | TSg | 32416 | Val20 attP2 |
| pnr | FBgn00031 | GATA4 | 30 | no | Hom_del | TSg | 34659 | Val20 attP2 |
| Pi3K21B | FBgn0020€ | PIK3R1 | 30 | snv_indel | Hom_del | TSg | 38991 | Val20 attP2 |
| Sox14; Sox | FBgn0005€ | SOX7 | 30 | snv_indel | Hom_del | TSg | 34794 | Val20 attP2 |
| park | FBgn00411 | PRKN | 28 | snv_indel | Hom_del | TSg | 38333 | Val20 attP2 |
| Dad | FBgn00204 | SMAD7 | 28 | snv_indel | Hom_del | TSg | 33759 | Val20 attP2 |
| Nedd4 | FBgn02591 | NEDD4L | 25 | snv_indel | Hom_del | TSg | 34741 | Val20 attP2 |
| Lkb1 | FBgn00381 | STK11 | 25 | snv_indel | Hom_del | TSg | 34362 | Val20 attP2 |
| nej | FBgn0261€ | CREBBP | 24 | snv_indel | Hom_del | Onc/TSg | 37489 | Val20 attP2 |
| ft | FBgn0001C | FAT4 | 23 | snv_indel | no | TSg | 34970 | Val20 attP2 |
| Phlpp | FBgn00327 | PHLPP1 | 23 | snv_indel | Hom_del | TSg | 57399 | Val20 attP40 |
| Utx | FBgn02607 | KDM6A | 22 | snv_indel | Hom_del | TSg | 34076 | Val20 attP2 |
| Bap170 | FBgn0042C | ARID2 | 21 | snv_indel | Hom_del | TSg | 26308 | Val10 attP2 |
| CG4887 | FBgn00313 | RBM10 | 21 | snv_indel | no | TSg | 65092 | Val20 attP40 |
| CG11505 | FBgn00354 | LARP4B | 19 | snv_indel | Hom_del | TSg | 64937 | Val20 attP40 |
| faf | FBgn0005€ | USP9X | 19 | snv_indel | Hom_del | TSg | 35728 | Val20 attP2 |
| unc-5 | FBgn0034C | UNC5D | 18 | snv_indel | Hom_del | TSg | 33756 | Val20 attP2 |
| Rbf; Rbf2 | FBgn00157 | RB1 | 15 | snv_indel | Hom_del | TSg | 65929 | Val20 attP2 |
| Set2 | FBgn00304 | SETD2 | 15 | snv_indel | Hom_del | TSg | 33706 | Val20 attP2 |
| Tet | FBgn02633 | TET1 | 15 | snv_indel | no | TSg | 62280 | Val20 attP40 |
| zfh1 | FBgn0004€ | ZEB1 | 15 | snv_indel | no | Onc/TSg | 29347; 6879 | Val10 attP2, and UAS-zfh1 |
| Chd1 | FBgn02507 | CHD1 | 14 | snv_indel | Hom_del | TSg | 34665 | Val20 attP2 |
| Nf1 | FBgn00152 | NF1 | 14 | snv_indel | Hom_del | TSg | 53322 | Val20 attP40 |
| ptc | FBgn00038 | PTCH1 | 14 | snv_indel | no | Onc/TSg | 28795 | Val10 attP2 |
| Eph | FBgn00259 | EPHA2 | 13 | snv_indel | no | Onc/TSg | 39066; 59844 | Val20 attP40, and UAS-Eph |
| Mkk4 | FBgn00243 | MAP2K4 | 13 | snv_indel | Hom_del | TSg | 42832 | Val20 attP40 |
| CG34404 | FBgn00854 | MCC | 13 | snv_indel | Hom_del | TSg | 67332 | Val20 attP40 |
| cic | FBgn02625 | CIC | 12 | snv_indel | no | TSg | 25995 | Val10 attP2 |
| fus | FBgn00234 | ESRP1 | 12 | snv_indel | no | TSg | 55921 | Val20 attP40 |
| Lerp | FBgn0051C | IGF2R | 12 | snv_indel | Hom_del | TSg | 57436 | Val20 attP40 |
| N | FBgn0004€ | NOTCH1 | 12 | snv_indel | no | Onc/TSg | already in our library | |
| XNP | FBgn00393 | ATRX | 11 | snv_indel | no | TSg | 32894 | Val20 attP2 |
| vap | FBgn00039 | RASA1 | 11 | snv_indel | Hom_del | TSg | 51793 | Val20 attP40 |
| SA; SA-2 | FBgn0020€ | STAG2 | 11 | snv_indel | no | TSg | 33395 | Val20 attP2 |
| Trpm | FBgn02651 | TRPM1 | 11 | snv_indel | Hom_del | TSg | 44503 | Val20 attP2 |
| aop | FBgn0000C | ETV6 | 10 | no | Hom_del | TSg | 34909 | Val20 attP2 |
| Mhcl | FBgn0026C | MYO18B | 10 | snv_indel | no | TSg | 51456 | Val20 attP40 |
| NetB | FBgn00157 | NTN1 | 10 | no | Hom_del | TSg | 34698 | Val20 attP2 |
| polybromo | FBgn00392 | PBRM1 | 10 | snv_indel | Hom_del | TSg | 32840 | Val20 attP2 |
| Ptp36E | FBgn02674 | PTPRK | 10 | snv_indel | Hom_del | TSg | 65919 | Val20 attP40 |
| Iswi | FBgn0011€ | SMARCA1 | 10 | snv_indel | no | TSg | 32845 | Val20 attP2 |

### GOF-Gain Of Function

| Fly_gene | FlyBaseID | Human_ge | Frequency | Mut_Type_ | Mut_Type_CNA | Role | Transgenic | Type transgenic |
|---|---|---|---|---|---|---|---|---|
| Pi3K92E | FBgn00152 | PIK3CA | 178 | snv/indel | high_level_amplification | Onc | 8287 | UAS-Pi3K92E |
| NA | NA | MYC | 65 | no | high_level_amplification | Onc | Already in our library | |
| egg | FBgn0086S | SETDB1 | 34 | no | high_level_amplification | Onc | 94136 | UAS-egg |
| arm | FBgn00001 | CTNNB1 | 32 | snv/indel | no | Onc | 8370 | UAS-arm |
| aurA | FBgn00001 | AURKA | 26 | no | high_level_amplification | Onc | 8376 | UAS-aurA |
| Egfr | FBgn00037 | ERBB2 | 26 | snv/indel | high_level_amplification | Onc | 9535 | UAS-Egfr |
| neb | FBgn00043 | KIF14 | 26 | no | high_level_amplification | Onc | 23707 | UAS-neb |
| salm | FBgn02616 | SALL4 | 25 | no | high_level_amplification | Onc | 29716 | UAS-salm |
| GATAd; pnr | FBgn00322 | GATA6 | 24 | no | high_level_amplification | Onc/TSg | 7223 | UAS-pnr |
| Galphas | FBgn00011 | GNAS | 24 | snv/indel | high_level_amplification | Onc | 6489 | UAS-Galphas |
| Akt1 | FBgn00103 | AKT3 | 23 | no | high_level_amplification | Onc/TSg | 8191 | UAS-Akt |
| Ptp61F | FBgn02674 | PTPN1 | 23 | no | high_level_amplification | Onc/TSg | 56194 | UAS-Ptp61F |
| tai | FBgn0041C | NCOA3 | 19 | no | high_level_amplification | Onc | 6378 | UAS-tai |
| Myb | FBgn00029 | MYBL2 | 17 | no | high_level_amplification | Onc | 83147 | UAS-Myb |
| pbl | FBgn0003C | ECT2 | 16 | no | high_level_amplification | Onc | 66160 | UAS-pbl |
| cnc | FBgn02629 | NFE2L2 | 16 | snv/indel | high_level_amplification | Onc/TSg | In Rusten stocks | |
| luna | FBgn00407 | KLF5 | 15 | no | high_level_amplification | Onc/TSg | 56814 | UAS-luna |
| Src64B | FBgn02627 | LYN | 15 | no | high_level_amplification | Onc | already in library | |
| CycE | FBgn00103 | CCNE1 | 14 | no | high_level_amplification | Onc | already in library | |
| Rok | FBgn00261 | ROCK1 | 14 | no | high_level_amplification | Onc | 6669 | UAS-Rok catalytic domain |
| SoxN | FBgn00291 | SOX2 | 14 | no | high_level_amplification | Onc | 83300 | UAS-SoxN |
| Src64B | FBgn02627 | SRC | 14 | no | high_level_amplification | Onc | already in library | |
| Pvf1 | FBgn00309 | VEGFA | 12 | no | high_level_amplification | Onc | 58426 | UAS-Pvf1 |
| Cdk4 | FBgn00161 | CDK4 | 11 | no | high_level_amplification | Onc/TSg | 6631 | UAS-Cdk4 |
| Tor | FBgn00217 | MTOR | 11 | snv/indel | no | Onc/TSg | Already in library | |
| Mitf | FBgn02631 | TFEB | 11 | no | high_level_amplification | Onc | Helene stocks HK01300 | |
| Skp2 | FBgn00372 | SKP2 | 10 | no | high_level_amplification | Onc | 59036 | UAS-Skp2 |
| SF2 | FBgn02834 | SRSF1 | 10 | no | high_level_amplification | Onc | 94907 | UAS-Sf2::GFP |
